# Theory of mind predicts conversational success in early right hemisphere stroke recovery

**DOI:** 10.64898/2026.04.23.720418

**Authors:** Andrea Suazo, Margaret Lehman Blake, Tatiana Schnur

**Author notes:** Corresponding to: **Tatiana Schnur,** NeuroRecovery Research Center, TIRR Memorial Hermann, 1333 Moursund St, Room 310, Houston, TX 77030, USA.

## Abstract

Those living with right hemisphere damage (RHD) often struggle with engaging in aspects of conversation that require understanding what a speaker means. There is growing evidence that conversation relies on deducing the speaker’s perspective, an ability known as theory of mind (ToM). However, whether individual differences in conversation after RHD relate to ToM deficits is unknown.

Here, we related individual differences in conversational success to ToM in 31 speakers during the early stages of RHD (median 5 days post-stroke) in comparison to 16 age- and education-matched controls. We measured conversational success as the number of differences identified between two images while participants conversed with study coordinators. A non-verbal false belief task measured ToM abilities. Baseline cognitive tasks assessed visual inattention, inhibitory control/visual selective attention and working memory.

Linear mixed-effects modeling revealed that the ToM ability to explicitly infer others’ perspectives while managing conflict with one’s own reliably predicted conversational success (β = 0.42, *p* = .023; β’s = .44 - .64 across sensitivity models). Adding false belief ToM substantially increased model fit (ΔR^2^ = .115, Cohen’s *f*^2^ = .15), independent of baseline cognitive abilities and demographic factors.

These findings provide the first empirical evidence in unilateral RHD that the ability to reason about a partner’s knowledge and manage conflict with one’s privileged perspective is critical for successful conversation. Results support theoretical models of ToM as a cognitive basis for everyday conversation. The clinical implications underscore the importance of socio-cognitive screening and ToM-based interventions to enhance communication outcomes in stroke rehabilitation.

## Introduction

After unilateral right hemisphere damage (RHD), over 50% of individuals experience consequential communication deficits which impair conversational ability [1–3]. Individuals with RHD may struggle to understand non-verbal cues such as intonation and facial expressions [4], fail to comprehend non-literal language and jokes [5–7], provide less relevant information [8], or mismanage conversational flow by speaking verbosely, asking fewer questions, taking turns inappropriately or failing to maintain topic [4,6,9,10]. Conversation is crucial for social participation, yet these RHD impairments are often missed by traditionally used language assessments in acute to subacute brain injury, like the Quick Aphasia Battery, which focus on structural language impairments rather than conversational and pragmatic skills [11]. Without more obvious signs of communication difficulties, RHD patients receive little to no speech-language intervention [12] and are discharged without addressing the impairments, which leads to challenges with social participation and potential social isolation [1]. Identifying the cognitive mechanisms that support conversation after RHD is therefore critical for understanding, testing for and treating these impairments.

One potential mechanism underlying these conversational deficits is the ability to understand that others have knowledge, beliefs and opinions that differ from one’s own (cognitive theory of mind; ToM) [13]. ToM is often impaired after RHD [14,15] (for reviews see Adams et al. [16]; Bomfim et al. [17]; Gupta et al. [18]). Individuals with RHD perform worse compared to healthy adults and individuals with left hemisphere damage on ToM tasks requiring mental state attribution, such as discerning a character’s false belief or inferring intentions in stories or videos [14,19–22]. Because successful conversation likely depends on speakers and listeners inferring each other’s mental states, and aligning and establishing a common ground, ToM deficits may directly contribute to conversational difficulties after RHD [23–31].

ToM comprises at least two partially dissociable operations that differ in how much they require managing competition between self and other perspectives [20,32–38]. These operations are often contrasted in terms of whether reasoning about another’s mental state is triggered spontaneously vs prompted by an explicit judgment. However, they also differ in the degree of self-perspective conflict they create. The first involves spontaneously inferring another person’s thoughts or perspective without being prompted, which typically entails relatively lower conflict with one’s self-perspective (here, other ToM). A second requires an explicit judgment about the other person’s perspective, increasing the need to resist interference from one’s own privileged perspective when it conflicts with the other person’s perspective (here, self ToM). Effective conversation is thought to require both knowing each other’s mental states to distinguish between one’s own privileged information vs. that which is shared and the resolving of perspective conflicts in real time [30,31,39]. Depending on conversation demands, either one or both of these aspects of ToM may support conversational success after RHD.

Despite growing recognition of the link between ToM and social communication, research directly connecting ToM to spoken conversation after RHD remains limited. Most of the evidence linking ToM and social communication comes from non-stroke populations. For example, in typically developing children, ToM skills support effective collaborative communication strategies such as giving sufficient information and requesting clarification [40]. In children with autism, ToM predicts the ability to maintain conversational topics [41]. Similarly, in neurotypical adults, stronger ToM is associated with fewer errors and thus greater success in a collaborative conversation task [42]. In monologic narrative discourse for aging adults, stronger ToM is linked to greater lexical informativeness and fewer coherence errors [43,44]. In work with individuals with chronic traumatic brain injury, ToM independently predicts pragmatic communication, such as appropriately producing or completing communicative exchanges, above and beyond executive function ability [45,46].

In the only study to date directly examining ToM and communicative production after RHD, people with ToM impairments used fewer appropriate referring expressions during monologic tasks (e.g. storytelling) to indicate to the listener to which character they were referring [19]. While suggestive, this work did not measure communicative success in interactive conversation. Critically, no studies examine this relationship in the early post-stroke period, when behavioral performance more closely reflects lesion effect prior to functional reorganization. Thus, the extent to which ToM supports successful conversation after RHD remains unknown.

Here, we tested whether individual differences in negotiating information with a conversation partner after RHD relate to individual differences in inferring another person’s perspective and managing conflict with one’s own perspective, both key aspects of ToM. This study offers several advances. First, we tested abilities in speakers during the early post-stroke period (median 5 days after stroke), capturing behavior before significant compensatory strategies or neural reorganization occurred. Second, we quantified conversational success in a naturalistic context using an interactive spot-the-difference task [47]. Third, we assessed ToM using a non-verbal false belief paradigm that dissociates different aspects of ToM, the spontaneous inference of another’s beliefs under relatively low self-perspective conflict (other ToM) and the explicit inference of another’s beliefs which increases conflict between one’s own and another’s perspective (self ToM) [48]. Importantly, this task minimizes linguistic and task-specific confounds between conditions like making judgements or sustaining attention across the task [49]. Lastly, we measured visual inattention, inhibitory control/visual selective attention and working memory to determine the unique contribution of ToM to successful conversation. We hypothesized that poorer ToM after RHD, particularly deficits in managing one’s privileged perspective with others’ (self ToM) would be associated with less successful conversation when negotiating conflicting information. To our knowledge, this is the first study to examine the relationship between individual differences in successful conversation and ToM abilities before reorganization of function in acute to subacute RHD after stroke.

## Materials and methods

### Participants

#### RHD Sample & Testing Timeline

We included 31 native monolingual English-speaking participants (17 male, 24 self-reported right-handed) with acute ischemic or parenchymal hemorrhagic right-hemisphere focal stroke (referred to as RHD) consecutively enrolled from inpatient stroke units at hospitals in the Texas Medical Center, in Houston, Texas. Included participants presented with lesions localized to the right cerebral hemisphere confirmed via clinical neuroimaging (CT or MRI) and electronic medical record (EMR) review. See Fig. 1 for the distribution of acute lesion damage in 25 participants with available clinical MRI (see Supplementary Material online for image acquisition and analysis details). We enrolled participants regardless of specific lesion location, stroke severity, or post-stroke communication status. This unselected sampling approach allowed for a heterogeneous cohort that captured a wide range of early post-stroke recovery. We completed behavioral testing acutely within an average of 3 days after stroke onset (n = 21 participants; SD = 3 days; range = 1–7 days) or subacutely, at an average of 40 days after stroke onset (n = 10 participants; SD = 9 days; range = 30–57 days). Overall, the median time from stroke onset to testing was 5 days (mean = 15 days, SD = 18 days, range = 1–57 days).

**Figure 1.**
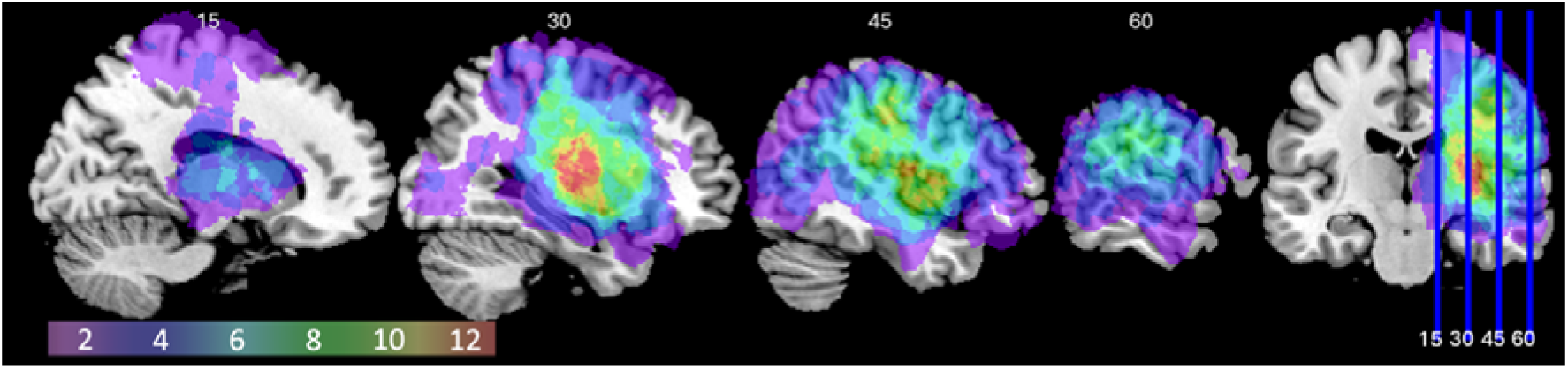
Acute lesion distribution across the right hemisphere (n = 25). Lesion overlap map displaying 4mm sagittal slices; (MNI *x* = 15, 30, 45, 60). Colors indicate the number of participants with a lesion at each voxel. Peak overlap occurred in subcortical regions (putamen/insula, caudate, pallidum) with additional overlap in peri-Rolandic frontal and parietal cortex and superior temporal regions (including Heschl’s gyrus). Mean lesion volume was 45.98 cm^3^ (range 0.25–322.11 cm^3^).

#### Screening Pipeline & Clinical Eligibility

As detailed in the participant flow diagram (Fig. 2), a total of 264 acute stroke admissions were screened for eligibility, of which 90 participants met inclusion criteria for RHD stroke. Per study protocol, exclusion criteria included if clinical assessment or EMR documentation indicated a prior left-hemisphere stroke, more than one prior right-sided stroke, history of other significant health conditions which could impact cognition or communication (e.g. dementia, schizophrenia, autism), or documented uncorrectable sensory deficits (i.e., blindness or deafness). Individuals were subsequently excluded prior to testing completion due to medical fragility (including severe visual inattention) or lack of alertness/cooperation, discharge prior to or interruption of assessment, or technical or equipment issues. We considered visual inattention severe when observed difficulty processing visual stimuli prevented participants from completing task requirements (e.g., inability to complete spatial attention tasks or to follow ToM task instructions after reporting seeing one of two displayed boxes). Decisions to discontinue visually based tasks reflected observed task performance, considered together with participant report and available clinical information. We tested but subsequently excluded two participants with predominantly right cerebellar strokes to focus the analyses on right cerebral hemisphere damage. We documented sensory and motor speech characteristics during acute hospitalization: eight participants self-reported a hearing difficulty, for which we implemented accommodations during testing (e.g., use of personal hearing aids or speaking directly to the participant’s preferred side), and 10 participants had a speech-language pathologist diagnosis of the presence of dysarthria in their medical records from the acute hospitalization. Two participants had a prior right hemisphere stroke. Because testing required a baseline level of alertness, cooperation, and visual/auditory processing sufficient to complete task instructions, our final analyzed cohort (n = 31) captured strokes in the minor, moderate, and moderate to severe range (admission NIH Stroke Scale Score mean = 7, SD = 5, range 0-17) [50]. Consequently, the sample preserved basic conversational ability at the aggregate group level while exhibiting broad individual variability across cognitive abilities.

**Figure 2.**
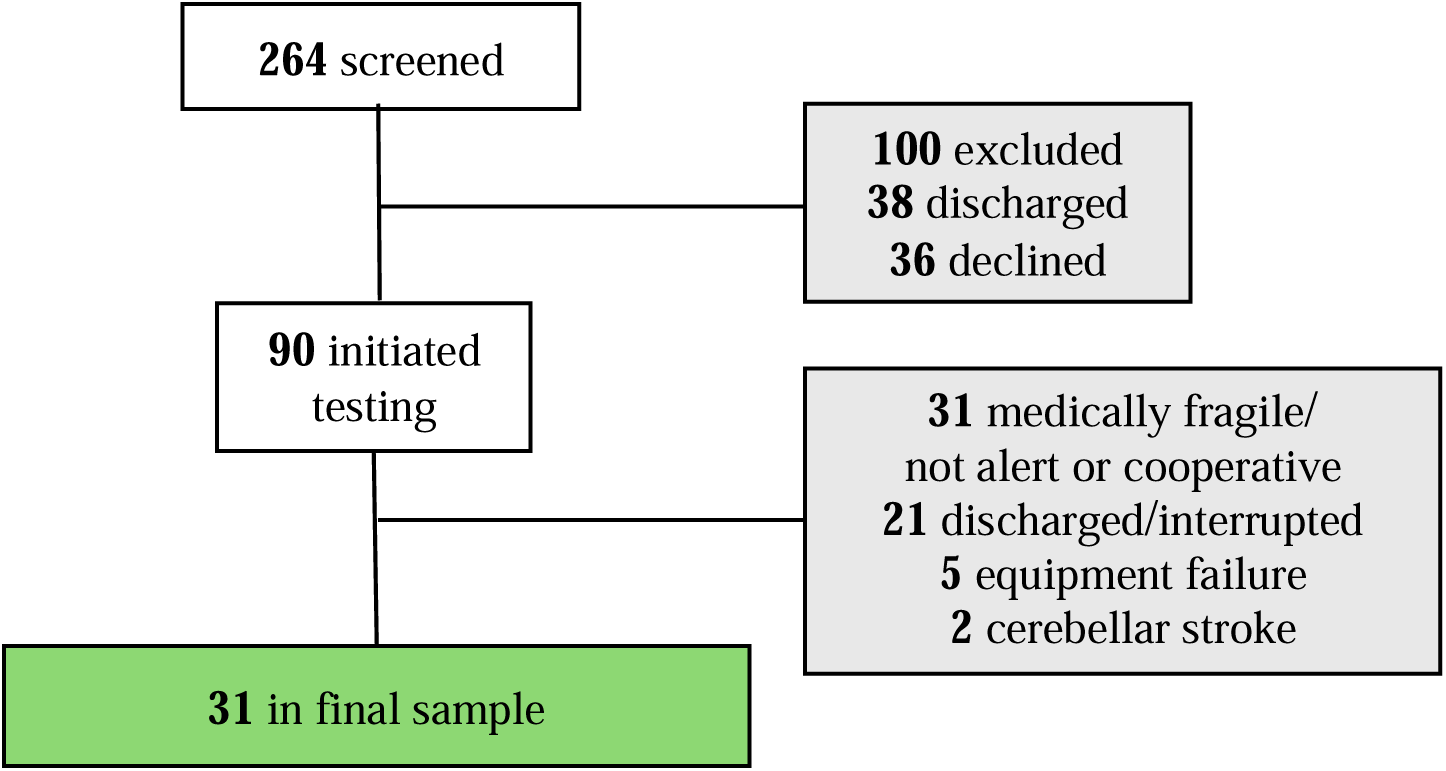
Participant enrollment and exclusion flowchart in early right-hemisphere damage (RHD). Flowchart detailing participant screening, exclusions, and final sample inclusion (n = 31). The final included sample is displayed in the green box; screening and exclusion stages are displayed in grey boxes.

#### Control Group Selection & Matching

For comparison with the RHD group, we initially recruited 21 neurotypical native English speakers from the surrounding community with no history of stroke, psychiatric or neurologic impairment, and normal or corrected-to-normal vision and hearing. We excluded five controls prior to analysis who either did not produce enough conversation (n = 1) or exhibited non-meaningful task compliance (scoring < 50%, n = 4). To verify normal cognitive status, control participants were screened for the potential presence of cognitive impairment using the Montreal Cognitive Assessment (MoCA) [51] with an alternative cut-off score of 23 to reduce the effect of bias [52–54]. The final control sample (n=16 participants; 7 male, 15 self-reported right-handed) had a mean MoCA score of 27 (SD = 2, range = 23–30), age of 54 years (SD = 15 years, range = 32–76 years) and 14 years of education (SD= 2 years, range 12–20 years). The RHD cohort (n = 31) had a mean age of 57 years (SD = 14 years; range = 30–81 years) and 13 years of education (SD = 2 years; range = 10–18 years). Welch’s *t*- and Chi-square tests confirmed that the RHD and Control groups did not significantly differ in age, education, or sex distribution (age *t*(30.9) = -0.82, *p* = .42; education *t*(26.6) = 1.11, *p* = .28; and sex *x^2^(1)* = 0.17, *p* = .68).

#### Ethical Approval & Informed Consent

Informed consent was obtained from either the participant or a legally authorized representative. The study was conducted under a single Institutional Review Board (sIRB) agreement, with UTHealth Houston and Baylor College of Medicine serving as the reviewing IRB (IRB of record) at different times during the study. All participating sites operated under the approved sIRB reliance arrangements. All methods were carried out in accordance with relevant guidelines and regulations.

### Procedure

The conversation, ToM tasks, and baseline cognitive tasks were administered and recorded as part of a ∼1.5 hour testing battery (Supplementary Fig. S1). Participants either completed the tasks in-person (n = 30 RHD participants, n = 10 controls) or remotely via Zoom (n = 1 RHD participants, n = 6 controls). Participants completed the conversation task with a research team member (AS, DB or KG, and in some cases for control participants EC or SM; see *Acknowledgements*).

#### Conversation Task

To elicit spontaneous and goal-oriented conversation between two partners, we administered a spot-the-difference task (Diapix) [47,55]. The Diapix has been shown to generate equal balance of speech between conversation pairs in non-clinical populations [47,55], and has been used to assess communication across different populations (i.e. dysarthric adults [56]; hearing-impaired adults [57]; healthy children/adolescents [58]). Conversation partners each received similar, standardized, difficulty-matched parallel picture sets [47] of either a beach (RHD n = 19, control n = 14), street (RHD n = 8, control n = 2), or farm/park scene (RHD n = 4, control n = 0). The picture sets differed in 12 ways, including the color of items, the wording on signs, or missing elements. Partners worked together without looking at each other’s pictures to find the differences. To successfully find differences, conversation partners both needed to contribute to the conversation and understand that the other held knowledge that they did not.

To mitigate participant differences in task strategy, we instructed participants to take the lead, to start at the top-right hand corner, and to move clockwise when describing the picture. We practiced the Diapix task with each participant for 3 to 5 minutes or until we found three differences with a training picture set. If a participant failed to understand task instructions and/or did not contribute during the training, we discontinued the task. After the practice, the researcher introduced the test picture set. The goal was to collect at least 4.5 minutes of conversation. For one control and one RHD conversation, the participant and researcher found the 12 differences in less than 4.5 minutes and thus, continued to a second picture set. To measure conversational success, we calculated the number of differences found by the conversation partners in 4.5 minutes (see Supplementary Material for transcription details). Because most participants only saw one picture set, conversational success scores were capped at 12.

#### Theory of Mind Task

We used a visual non-verbal false belief task to assess two aspects of ToM which differentially assessed the perspective that a viewer must take to understand another person’s beliefs [48]. The first component, other ToM, assessed the ability to spontaneously infer another person’s perspective without explicit instruction to do so, under relatively low self-perspective conflict. The second component, self ToM, assessed the explicit ability to infer another person’s perspective while managing conflict between self-perspective and another person’s perspective. Similar to classic ToM tasks (for a review see Martín-Rodríguez and León-Carrión [59]), the Biervoye et al.’s [48] assessment used scenarios in which the viewer had more information than a character in the scene and had to decide what the character knew. Participants saw a series of short video trials with no sound, with the following structure. A man placed a green object in one of two opaque boxes in sight of a woman. She left the room and then the man either kept the boxes in the same position or switched them. Conditions differed across trials. In all false belief trials (whether other or self ToM) the man switched the location of the green object. However, in the other ToM false belief condition, the participant did not see where the object was initially placed. When the woman returned to the room, she indicated where the object was with a pink marker. The participant then answered the question “show me where the green object is”. For correct other ToM false belief trial responses, the participant had to understand that the woman had a false belief about the object location because the boxes were switched during her absence. As a result, the woman put the pink marker on the wrong box and the green object must have been in the other box. Since the participant did not see the object’s location throughout the video, participants did not have especially strong self-perspective competing knowledge about the location. However, correct responding still required an indirect inference (discounting the woman’s cue), so this condition was not a process-pure measure of automaticity. In contrast, in the self ToM false belief trials the participant saw where the object was hidden initially. When the woman returned, she did not indicate where the object was. Instead, participants answered the question “Which box will she open first to find the object?”. In this condition, the participant already knew the location of the object but to correctly respond they must have understood that the woman had a false belief because she did not see the switch. Thus, when asked where the woman will look for the object, they had to manage conflict between their knowledge and the woman’s false belief.

Interleaved control trials assessed the within-task contributions of working memory, attention and comprehension, independent of ToM ability while controlling for unilateral neglect. In the other ToM condition, memory control trials assessed working memory where the woman indicated the object location before the object location switched. This required the participant to remember the object location and update the location when it switched at the end of the trial but did not require inference about the woman’s knowledge. In other ToM filler control trials, the woman indicated where the object was before leaving the room, and then the man removed the object from one box and placed it in the other box, giving the participant a visual confirmation of the object switch. Therefore, the participant did not need to infer the woman’s knowledge to respond, just an understanding of task instructions. Filler trials also confirmed that the woman was genuinely trying to assist in identifying the object location. In self ToM filler control trials, the participant saw that the man switched the object’s location in the presence of the woman. Because there was no conflicting knowledge between the object location and the woman’s knowledge of the object location, the participant only needed an understanding of task instructions to respond. Poor performance on filler trials demonstrated a deficit in some aspect of input processing beyond any potential deficit in ToM reasoning. Lastly, in the true belief control trials (which also served as memory control trials for the self ToM condition) any manipulation of the object (e.g. lifting a box rather than switching boxes) never changed the object’s location. If a participant made more errors on the true belief trials (when the location never changed) compared to the false belief trials, this suggested that the participant used a non-ToM based strategy in the false belief trials such as picking the box which the woman did not point at. Because control trial types were designed to assess non-ToM processing demands of the task, we created a composite control score by averaging performance on working memory, true belief and filler control trial types for each ToM condition, respectively. See Table 1 for a summary of trial types.

**Table 1.**
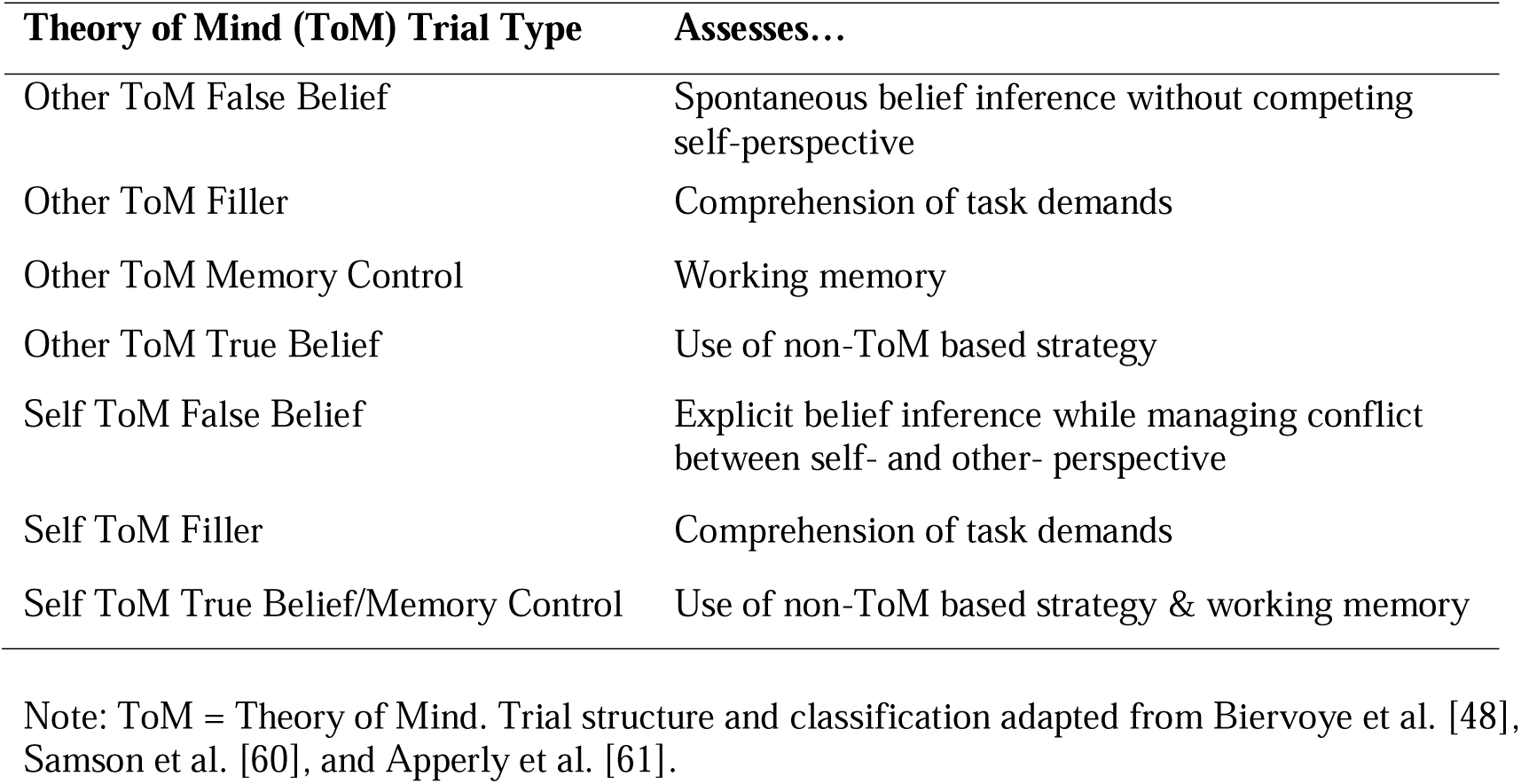
Summary of Theory of Mind (ToM) trial types and targeted constructs.

Following Biervoye et al. [48], we presented the other ToM condition before the self ToM condition to keep the demands of the other ToM condition high (also see Speiger et al. [62]). Participants received feedback after every trial. Before both ToM conditions, participants received task instructions and performed 2–4 practice filler trials. The experimental task was administered only if participants were accurate on two consecutive practice trials. We assessed ToM performance with four dependent variables: other ToM false belief trial accuracy (out of 8), other ToM control trial accuracy (out of 16), self ToM false belief trial accuracy (out of 8) and self ToM control trial accuracy (out of 16).

### Baseline Cognitive Tasks

#### Visual Inattention

The object (apples) cancellation task [63] and an adaptation of Ogden’s scene copy task [64] measured visual inattention. Both tasks assessed egocentric (viewer-centered) and allocentric (stimulus-centered) inattention. We averaged the accuracy score for the egocentric and allocentric inattention for each task to create a total visual inattention score for the object cancellation and scene copy tasks, respectively. The task accuracy scores were significantly, positively correlated (r = .65, p < .001). Thus, we averaged task scores to create a composite visual inattention accuracy score.

#### Inhibitory Control and Visual Selective Attention

The NIH Toolbox Flanker Inhibitory Control and Attention Test measured the capacity to inhibit automatic responses that interfere with achieving a goal, while also measuring selective attention (V1.28) [65]. Participants saw a row of five arrows on the screen and pressed the button that matched the direction that the middle arrow was pointing. The middle arrow pointed in the same direction as the flanking arrows in 12 congruent trials and pointed in the opposite direction as the flanking arrows in eight incongruent trials. The dependent measure was the NIH Toolbox age-, race-, gender-, and education-corrected T-score (normative sample mean = 50, SD = 10 [66]).

#### Working Memory

The digit span subtest from the Repeatable Battery for the Assessment of Neuropsychological Status assessed working memory [67]. Participants heard sequences of two to nine numbers at approximately one word per second, after which they were asked to repeat the sequence back in the correct order. There were two sequences per list length. The second sequence was administered only when participants failed the first one. We discontinued the task after failure of both sequences in any list length. Participants received an item raw score of two if they passed the first string in an item, one if they passed the second string and zero if they failed both. Each item’s score was summed for a total raw score range of 0 –16. See Supplementary Fig. S2 for performance distributions across baseline cognitive tasks and Supplementary Table S1 for summary statistics.

### Baseline expressive and receptive lexical-semantic access

For insight into the RHD sample’s language skills post stroke, we tested participants’ expressive and receptive lexical-semantic ability.

#### Word-Picture Matching

We administered an adapted word-picture matching task from Schnur & Lei [68] (cf. Breese & Hillis [69]). We presented one of 17 pictures to participants and asked, “Is this a ?” Participants responded “yes” if the picture matched the spoken word (e.g., CAT/cat) and “no” if they did not match. For unmatched pairs, 17 pictures were presented once with a semantic foil (e.g., CAT/dog) and another time with a phonologically related foil (e.g., CAT/hat). A total of 51 trials was divided into three presentation sets in a larger test battery. We calculated accuracy out of 51 trials.

#### Picture Naming

Participants named 30 pictures of depictable objects from the Philadelphia Naming Test-Short form [70]. Participants were shown each picture on a computer screen and asked to use one word to name the object with a naming deadline of 10s. Two research assistants supervised by a speech-language pathologist served as scorers. The single-measurement, absolute agreement, two-way mixed effects model ICC for error detection was .91 [.91, .93], indicating excellent reliability [71] for the two scorers. We calculated accuracy out of 30. See Supplementary Fig. S3 for performance distributions across lexical-semantic access tasks and Supplementary Table S1 for summary statistics.

### Statistical Analysis

For an intuitive interpretation of degree of impairment and for direct comparison between dependent measures, we *z*-scored conversational success accuracy, ToM accuracy, digit span raw score, word-picture matching accuracy and picture naming accuracy to control participant performance. We *z*-scored visual inattention composite scores to the RHD participant group. We transformed the automatically generated Flanker *T*-scores, which were normed to an age-, gender-, and education-matched control population [66], into *z*-scores.

We tested for outliers using a studentized residual criterion (> 2.5 SDs from the mean) and Cook’s distance (> 3 SD above the sample mean) [72]. One participant met exclusion criteria, leaving 30 RHD participants for the regression analyses and associated correlation matrices. Group summaries and task-performance comparisons included all 31 RHD participants.

We used hierarchical linear mixed-effects modeling to test whether ToM performance predicted conversational success, while controlling for the contribution of other cognitive abilities. To directly test our hypothesis whether ToM performance, specifically self ToM, predicted conversational success while accounting for non-ToM task demands, the primary model included the other-and self-ToM false belief measures and their corresponding ToM control measures. We then conducted three sensitivity analyses to examine whether this association was dependent on adjustment for other participant characteristics, adding: (1) baseline cognitive performance (visual inattention, inhibitory control/visual selective attention, working memory), (2) demographic/clinical variables (age, years of education, days post-stroke onset), and (3) both sets of covariates. In an additional targeted sensitivity analysis, we examined whether the association between self ToM and conversational success was robust to adjustment for lesion volume. Because lesion-volume data were available for a subset of participants, we repeated the primary model in this subset before and after adding lesion volume as a covariate (see Supplementary Material). Because participants completed the conversation task with one of three research team members, we included the conversation partner as a random intercept in all models. We omitted measures of lexical-semantic access as model covariates for three reasons: (1) overall high accuracy across the RHD cohort (average accuracy 94%; Supplementary Table S1 and Supplementary Fig. S3), (2) the non-verbal nature of the ToM task, which was specifically designed to minimize linguistic demands, and (3) preserving model parsimony to prevent overparameterization.

To assess overall model significance, we evaluated each model against an intercept-only null model using likelihood ratio tests. To evaluate the effect size of false-belief ToM in our models, we compared the hypothesis-driven primary model (containing ToM false belief + matched ToM control measures) to a reduced baseline model containing only the ToM control measures.

## Results

Pairwise correlations across the RHD hierarchical regression sample demographics and task performance are shown in Fig. 3 (see Supplementary Fig. S4 for an expanded correlation matrix including the baseline language tasks).

**Figure 3.**
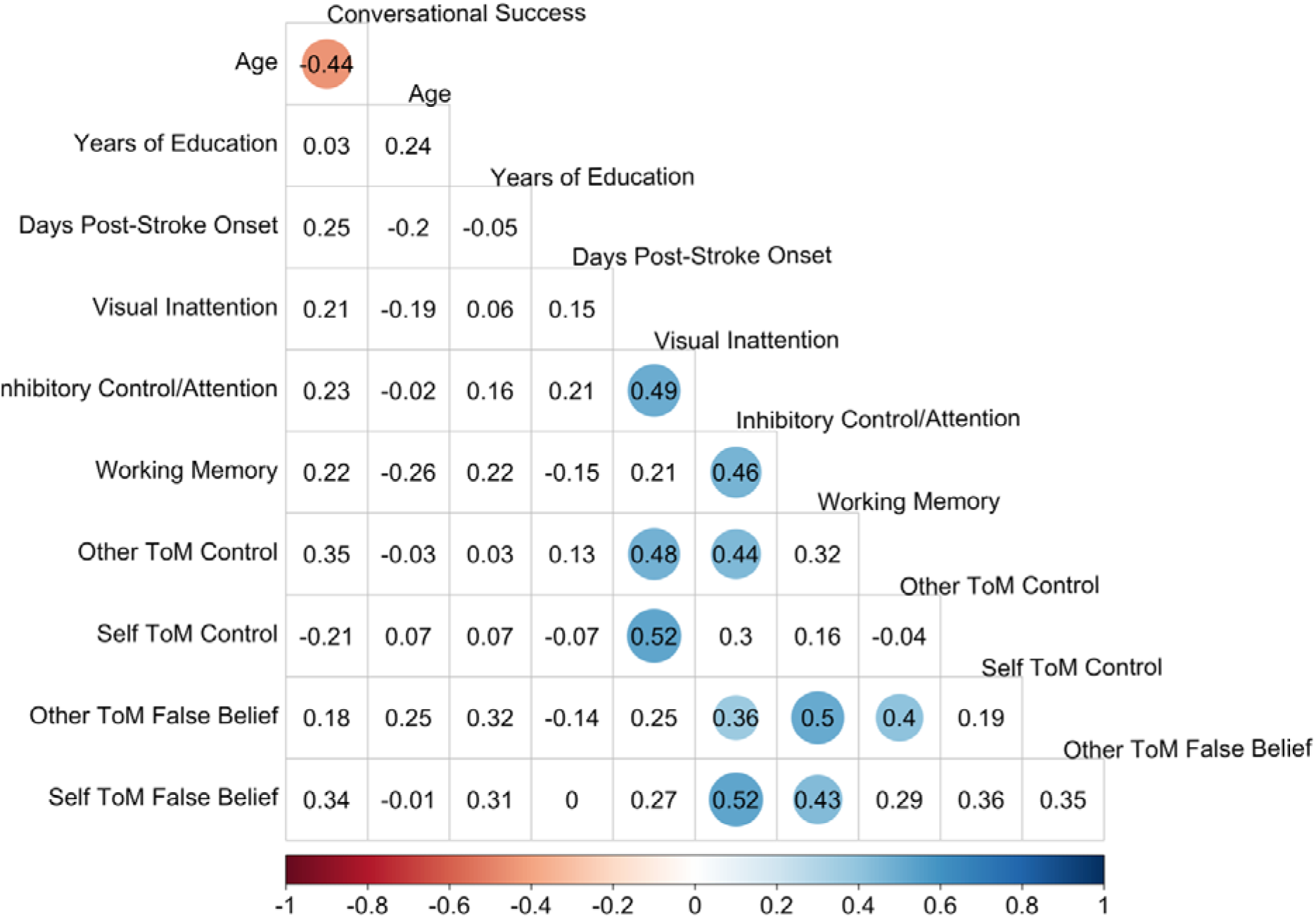
Correlations between demographic and clinical information, baseline cognition, Theory of Mind (ToM) performance and conversational success in early post-stroke right-hemisphere damage (RHD) (n=30). Matrix cells report pairwise Pearson’s *r* correlation values. Colored cells indicate significant correlations (*p* < .05). Circle size and color intensity indicate the magnitude and direction of the association (see color bar). RHD = right-hemisphere damage; ToM = Theory of Mind.

### ToM performance

On average across ToM conditions, RHD participants achieved 60% accuracy on false belief trials (range 0% – 100%) and 88% accuracy on control trials (range 69% – 100%), while controls were near ceiling on both (false belief trials range = 88% -100%; controls trials range = 88% - 100%). At the group level, RHD and controls significantly differed in false belief (38% difference, Welch’s *t*(32.1) = 7.55, *p* < .001) and control trial performance (11% difference, Welch’s *t*(35.3) = 6.67, *p* < .001). For RHD participants, other ToM false belief accuracy was lower and more variable than self ToM false belief accuracy (22% difference, paired *t*(30) = -3.22, *p* = .003), suggesting that the reduced self-perspective conflict in other ToM did not translate into lower overall difficulty in this paradigm. See Supplementary Fig. S5 for the ToM condition and trial performance breakdown across groups.

### Conversation (Diapix) performance

At the group level, conversational success was comparable between RHD (mean = 65%, SD = 18%) and neurotypical control participants (means = 69%, SD = 16%; Welch’s *t*(34.6) < 1, *p* = .51). However, performance within the RHD group exhibited substantial individual variability (range 33%–100%), establishing a wide continuum of conversational success across participants (Supplementary Fig. S6). Task instructions asked participants to take the lead in the conversation, to encourage participants to actively contribute to the conversation. Although control participants contributed a significantly higher proportion of total dialogue compared to RHD participants (mean = 59%, SD = 10%, range = 41% –77%; Welch’s *t*(35.8) = 3.50, *p* = .001), RHD participants maintained an average 47% of total dialogue words (SD = 12%, range = 26% –70%), which aligned closely with previous work [47]. This substantial verbal contribution, paired with group-level success in identifying picture differences, indicated that participants successfully engaged with the basic demands and goal of the conversation task. For the 10 participants with documented dysarthria, their participation (mean word count = 332 words, range =172 – 444 words) and success (average success = 65%, range 33% - 92%) was similar to those without dysarthria (n= 21; word count Welch’s *t*(27.4) < 1, *p* = .72; mean word count = 347 words, range 134 – 578 words; success Welch’s *t*(14.0) < 1, *p* = .95; average success = 65%, range 33% - 100%).

### Relating ToM and Conversational Success

The hypothesis-driven primary model (comprising self and other false belief ToM and associated control trial composites) accounted for 22% of the variance in conversational success (marginal R² = .218), with the random effect of conversation partner explaining an additional 25% of the variance (conditional R² = .458). Consistent with predictions, better self ToM false belief performance significantly predicted higher conversational success (self ToM false belief trials β =0.42, *p* = .023; see Fig. 4A and Table 2, Primary Model) whereas other ToM false belief performance was not a significant predictor (β =0.06, *p* =.734). Unexpectedly, worse performance on self ToM control trials predicted better conversational success (β = -0.41, *p* = .018).

**Figure 4.**
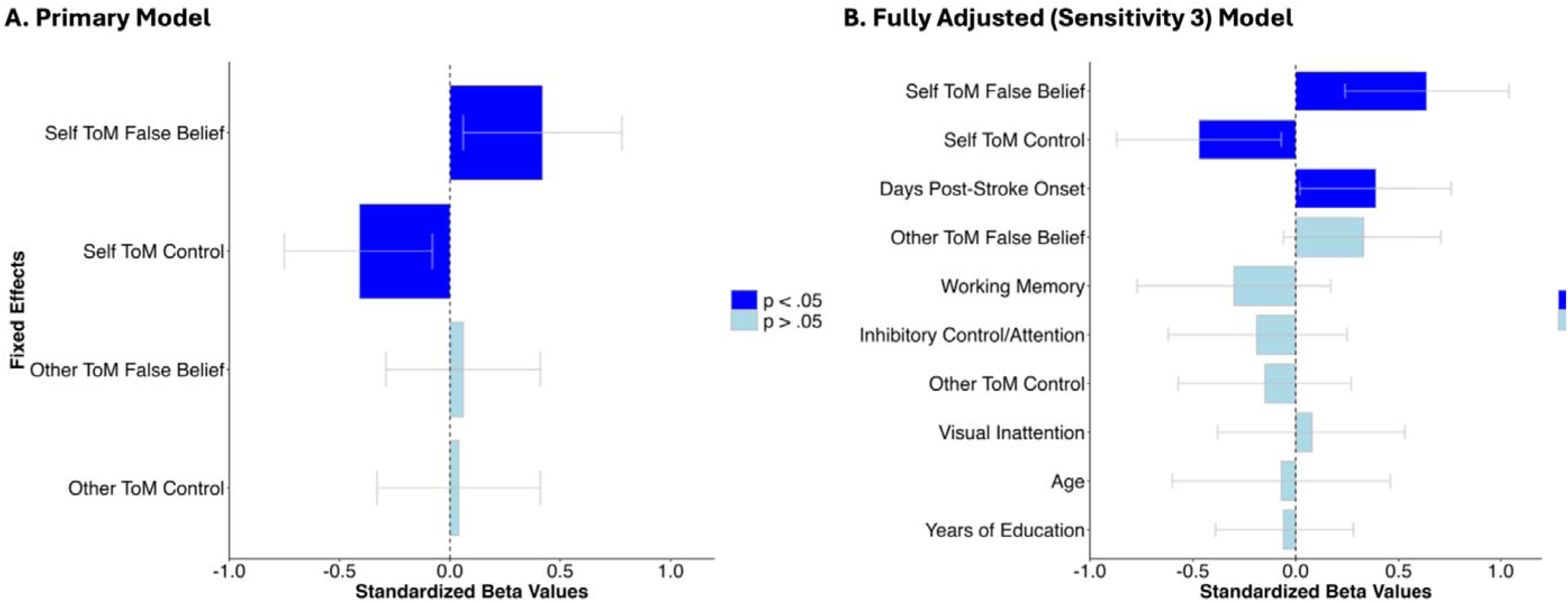
Theory of Mind (ToM) performance predicting conversational success across primary and fully adjusted (Sensitivity 3) models in early post-stroke right-hemisphere damage (RHD). Standardized fixed-effect estimates (β) with 95% confidence intervals from regression models predicting conversational success. (A) Primary model including core Theory of Mind (ToM) predictor composites. (B) Fully adjusted (Sensitivity 3) model incorporating ToM predictors alongside baseline demographic, clinical and cognitive covariates. Dark blue bars denote statistically significant predictors (*p* < .05); light blue bars denote non-significant predictors (*p* >= .05). Predictors are ordered by standardized beta value. RHD = right-hemisphere damage; ToM = Theory of Mind.

**Table 2.** Linear mixed-effects models predicting conversational success in early post-stroke right-hemisphere damage (RHD) (n=30).

| Fixed Effects | Primary Model<br>Core ToM |  |  | Sensitivity 1<br>Cognition Adjusted |  |  | Sensitivity 2<br>Demographic/Clinical<br>Adjusted |  |  | Sensitivity 3<br>Fully Adjusted |  |  |
| --- | --- | --- | --- | --- | --- | --- | --- | --- | --- | --- | --- | --- |
| | std. $\beta$ | std. 95% CI | *p* | std. $\beta$ | std. 95% CI | *p* | std. $\beta$ | std. 95% CI | *p* | std. $\beta$ | std. 95% CI | *p* |
| Other ToM Control | 0.04 | -0.33, 0.41 | .832 | -0.08 | -0.53, 0.36 | .698 | -0.09 | -0.47, 0.29 | .617 | -0.15 | -0.57, 0.27 | .466 |
| Self ToM Control | -0.41 | -0.75, -0.08 | .018* | -0.54 | -0.97, -0.12 | .015* | -0.42 | -0.75, -0.10 | .013* | -0.47 | -0.87, -0.07 | .024* |
| Other ToM False Belief | 0.06 | -0.29, 0.41 | .734 | 0.17 | -0.22, 0.56 | .381 | 0.19 | -0.17, 0.54 | .280 | 0.33 | -0.06, 0.71 | .094 |
| Self ToM False Belief | 0.42 | 0.06, 0.78 | .023* | 0.56 | 0.14, 0.99 | .011* | 0.44 | 0.09, 0.80 | .017* | 0.64 | 0.24, 1.04 | .004* |
| Visual Inattention |  |  |  | 0.21 | -0.28, 0.70 | .393 |  |  |  | 0.08 | -0.38, 0.53 | .731 |
| Inhibitory Control/Attention |  |  |  | -0.06 | -0.48, 0.36 | .773 |  |  |  | -0.19 | -0.62, 0.25 | .381 |
| Working Memory |  |  |  | -0.25 | -0.71, 0.20 | .258 |  |  |  | -0.30 | -0.77, 0.17 | .198 |
| Age |  |  |  |  |  |  | -0.07 | -0.52, 0.37 | .735 | -0.07 | -0.60, 0.46 | .796 |
| Years of Education |  |  |  |  |  |  | -0.08 | -0.41, 0.25 | .631 | -0.06 | -0.39, 0.28 | .733 |
| Days Post-Stroke Onset |  |  |  |  |  |  | 0.34 | 0.00, 0.67 | .047* | 0.39 | 0.02, 0.76 | .042* |
| Random Effects | RV ($\sigma^2$) | RIV ($\tau_{00}$) | ICC | RV ($\sigma^2$) | RIV ($\tau_{00}$) | ICC | RV ($\sigma^2$) | RIV ($\tau_{00}$) | ICC | RV ($\sigma^2$) | RIV ($\tau_{00}$) | ICC |
| Conversation Partner (n=3) | 0.78 | 0.34 | 0.31 | 0.78 | 0.50 | 0.39 | 0.66 | 0.47 | 0.41 | 0.61 | 0.85 | 0.58 |

Note: \* $p < .05$ . std. = standardized; CI = confidence interval; $p$ = $p$ -value; RV ( $\sigma^2$ ) = residual variance; RIV ( $\tau_{00}$ ) = random intercept variance; ICC = intraclass correlation; RHD = right-hemisphere damage; ToM = Theory of Mind.

To evaluate overall model significance, we compared the primary model to a reduced model containing only the ToM control measures. Adding the false belief ToM measures significantly improved model fit (χ²(2) = 6.66, *p* = 0.036) and increased marginal R² from .103 to .218 (ΔR² = .115). This 12% increase in unique variance corresponded to a moderate effect size (Cohen’s f^2^ = .15; [73])

To evaluate whether the association between ToM performance and conversational success was independent of participant characteristics, we conducted three sensitivity analyses adjusting for: baseline cognitive measures (Sensitivity 1), demographic and clinical variables (Sensitivity 2), and both sets of covariates in a fully adjusted model (Sensitivity 3; Table 2, Fig. 4B, and Supplementary Fig. S7). Across all sensitivity analyses, the positive relationship between self ToM false belief performance and conversational success remained consistent in magnitude (βs = .44 to .64). Specifically, this association remained statistically significant in the cognition-adjusted model (β = .56, *p* = .011), the demographic/clinical-adjusted model (β = .44, *p* = .017), and the fully adjusted model (β = .64, *p* = .004). Additionally, days post-stroke onset was positively associated with conversational success in the demographic-adjusted (p = .047) and fully adjusted (p = .042) models. Across sensitivity models, the relationship between other ToM false belief performance and conversational success remained non-significant (*p*’s ≥ .094). Of note, adjustment for lesion volume in the subset of participants with available data (N = 24) did not substantially alter the magnitude or direction of the estimated self ToM association with conversational success (see Supplementary Table S3).

To assess whether the predictors explained additional variance in conversational success, all models were evaluated against an intercept-only null model. All except the cognition- adjusted model (p =.052) provided significantly better fit than the null model (all other *p*’s ≤ .026; see Table 3 for complete Model Fit statistics). To assess multicollinearity (see Fig. 3), we calculated variance inflation factors (VIFs) for the fully adjusted model. All VIFs were < 2.4, indicating low collinearity among predictors [74]. Adjusted partial regression scatterplots illustrate these relationships after adjusting for all full model covariates (Supplementary Fig. S8). freedom; R^2^_m_ = marginal; R^2^ (variance explained by fixed effects), R^2^_c_ = conditional R^2^ (variance explained by fixed and random effects).

**Table 3.**
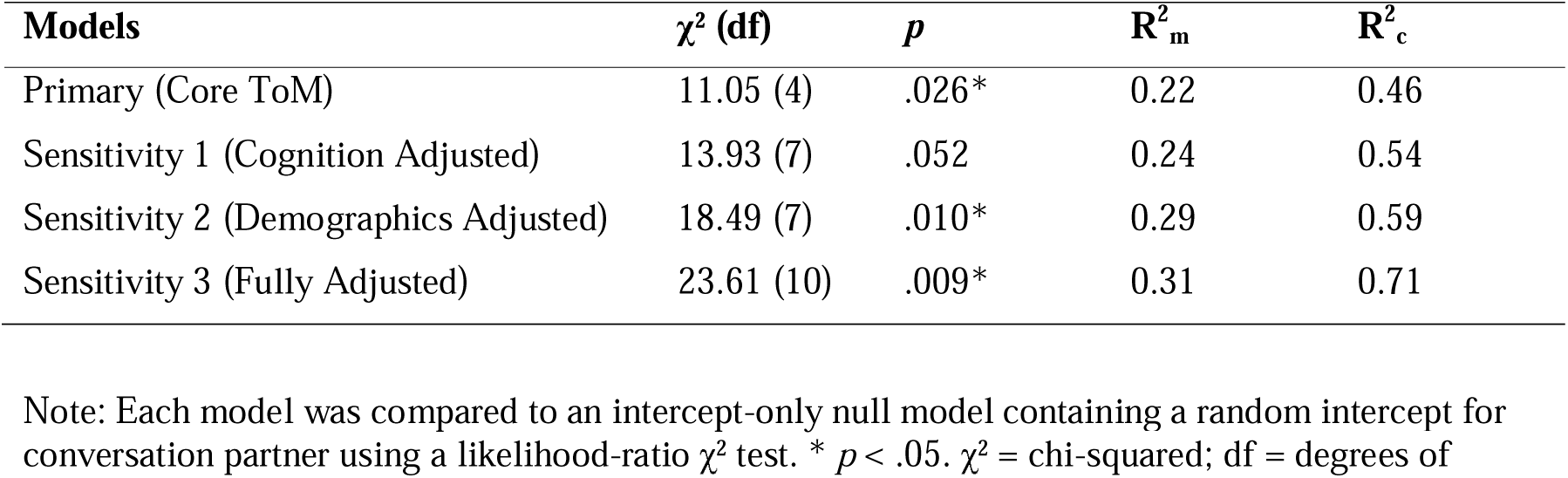
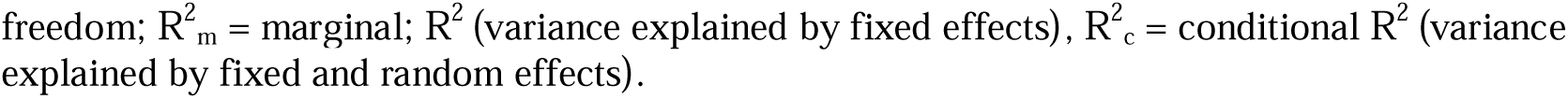
Linear mixed effects models fit statistics.

The negative association between self ToM control performance and conversational success remained consistent across all models (βs = -.41 to -.54, *p*’s = .013 - .024). To better clarify this unexpected relationship, we conducted an exploratory follow-up analysis in the primary model by decomposing the self and other ToM control composite scores into their constituent trial types (filler, memory control (MC), and true belief (TB) trials; MC and TB combined as a single MC/TB condition for self ToM). In this decomposed model, self ToM false belief performance remained a significant predictor of conversational success (β = 0.46, *p* = .027), whereas the negative association previously observed for the control composite score was attenuated, with no individual self ToM control trial reaching statistical significance (all *p* ‘s> .203; See Supplementary Fig. S9 and Table S2). Taken together, these analyses indicated that the ability to explicitly infer a partner’s perspective while managing conflict with one’s own (self ToM false belief performance) reliably predicted conversational success, whereas any association involving self ToM control trials was non-robust and did not isolate to a specific task-processing component.

## Discussion

After right hemisphere damage (RHD), individuals often experience spoken communication impairments which reduce quality of life [1–3]. Successful collaborative conversation requires interlocutors to track both shared and privileged information while resolving conflicts between one’s own perspective and another’s [23–31]. Here, we investigated whether individual differences in conversational success during early recovery of stroke-related RHD were associated with differences in the socio-cognitive ability to infer others’ beliefs (theory of mind; ToM). During a collaborative conversation task, individuals with RHD achieved greater conversational success when they performed better on the self-perspective conflict ToM condition (self ToM), which places greater relative demands on managing interference from one’s own privileged perspective while reasoning about a partner’s perspective. This association was present over and above performance on the other perspective ToM condition (other ToM), which also requires belief reasoning but under relatively lower perspective conflict. The association held after accounting for demographic factors and broader cognitive abilities, indicating that self ToM contributes unique explanatory power for conversational outcomes in early RHD.

To our knowledge, this is the first evidence that individual differences in ToM explain unique variance in the success of spoken communication after RHD. These findings extend converging evidence that the right hemisphere supports ToM and communicative effectiveness [19,75,76] and complement work showing that false belief reasoning is engaged during conversation [29,77–79]. By focusing on dialogue rather than monologic production, the present study highlights ToM’s relevance to the dynamic, bidirectional demands of real time interaction.

### Managing Perspectives is Critical for Collaborative Conversation

The present pattern demonstrates that during collaborative conversation, conversational success following RHD is linked to the ability to manage conflict between one’s own privileged perspective and a partner’s perspective. While ToM is not the sole cognitive process supporting real-time interaction, its unique contribution is both theoretically and practically meaningful. Unscripted, naturalistic dialogue is an inherently complex behavior shaped by multiple factors (e.g., linguistic, dyadic etc.). In this context, incorporating false belief ToM measures alongside matched non-ToM task controls substantially improved our ability to account for communicative success. This gain highlights the unique variance in conversational success explained by false belief reasoning beyond baseline task demands, representing a meaningful effect. Ultimately, these findings show that perspective management directly supports natural dialogue.

In the Diapix conversation task [47], participants needed to determine what they knew, what their partner likely knew and how those perspectives differed, and then use that comparison to guide their utterances to accomplish the conversation goals. Meanwhile, both ToM conditions required understanding another person’s belief, but the self ToM condition placed greater demands on comparing that belief with one’s own privileged perspective and resisting interference from self knowledge, and the latter, when impaired, negatively impacted collaborative conversation. In contrast, other ToM performance was not significantly associated with conversation success. This null finding is unlikely to reflect measurement failure or floor/ceiling effects. Other ToM imposed substantial inferential demands with participants performing lower on this condition overall, and performance showed meaningful associations with education and working memory, indicating that the measure was sensitive to individual differences. Instead, this behavioral dissociation suggests that simply representing a partner’s perspective is insufficient to drive conversational success. Rather, the bottleneck in collaborative interaction is managing conflict between self and other perspectives during real-time perspective comparison. This interpretation of the conversation demands in this task is consistent with Heller and Brown-Schmidt’s Multiple Perspectives Theory [30,31], which proposes that communication depends on maintaining separate representations of self and other and on a comparison process that computes similarities and differences between them. The Diapix task places continuous pressure on this comparison process, where successful performance depends not simply on recognizing that a partner has a different perspective, but on using that asymmetry to decide what to say next.

The present findings also motivate a hypothesis regarding the differential involvement of other and self ToM for communication. Other ToM may contribute relatively more than self-perspective management in communicative situations where the central challenge is not resolving direct interference from one’s own privileged information, but instead inferring a speaker’s intended meaning from shared context and the speaker’s private mental states. Candidate cases include irony, deceit and other forms of non-literal language comprehension. For example, prior work suggests that understanding irony and deceit requires detecting a mismatch between a speaker’s utterances and the speaker’s private knowledge, where the listener must also track what is or is not shared between interlocutors [80]. This does not mean that such phenomena eliminate comparison between perspectives. Instead, compared with collaborative referential tasks such as Diapix [47], they may place less weight on resisting interference from one’s own privileged perspective and greater weight on deriving speaker meaning from the other person’s perspective and the shared context. On this view, belief reasoning under relatively lower self-perspective conflict (other ToM) may be more important for some forms of pragmatic comprehension, whereas belief reasoning under stronger perspective conflict (self ToM) may be particularly consequential for cooperative, goal-directed dialogue.

More broadly, the present findings align with evidence from clinical and non-clinical populations across the lifespan, showing that stronger ToM skills are related to better conversational outcomes [40–46]. Particularly relevant is work by Markiewicz et al. [42], where lower ToM competency led to increased communication errors in a cooperative conversation task in 350 healthy adults, supporting the idea that lower ToM increases opportunities for misinterpretation and communication breakdowns. Within this framework, we propose that impaired ToM after RHD contributes to poorer coordination of shared and privileged information, leading to poorer conversational success. Importantly, because we assessed individuals in the acute to subacute post-stroke window (median 5 days), these results likely reflect disrupted mechanisms before neural reorganization and compensatory strategies substantially reshape behavior. This timing strengthens the interpretation that early RHD disrupts ToM-related processes, that in turn, matter for functional communication.

### Alternative Explanations

ToM closely interacts with executive function across development and clinical populations, raising the possibility that apparent ToM effects on communication reflect broader executive limitations [32,81–91]. Executive abilities also relate to communication outcomes [92–96]. From this view, the self-perspective conflict condition may appear more associated with conversational success simply because it is more attentionally or more working-memory demanding than the other perspective ToM condition.

Several aspects of our data argue against a purely domain-general executive explanation. First, regression modeling demonstrated that self ToM accounted for unique variance in conversational success beyond baseline cognitive and demographic controls, indicating that self-perspective conflict management represents a distinct socio-cognitive mechanism rather than a proxy for broad executive control. Although self ToM correlated with inhibitory control/visual selective attention (NIH Toolbox Flanker) [65] and verbal working memory (digit span) [67], neither executive measure significantly predicted conversational success in our regression models. While these null associations should be interpreted cautiously given our sample size, they demonstrate that variation in general inhibition or working memory alone does not explain the observed relationship.

Second, task difficulty considerations further argue against a domain-general cognitive load explanation. Participants performed significantly worse on the other ToM false-belief condition than the self ToM condition. While self ToM requires overriding privileged self-perspective knowledge, other ToM requires a distinct, multiple step inference process. Participants must figure out where the woman believes the object is based on her incorrect cue, while simultaneously inhibiting the prepotent urge to select the box she indicated. That other ToM involved greater inferential and inhibitory demands, yet failed to associate with conversational success, reinforces that general cognitive load or cue inhibition does not account for the results. Instead, this behavioral dissociation suggests that the capacity to manage conflict between one’s privilege perspective and another’s serves as a key socio-cognitive mechanism supporting collaborative dialogue following right hemisphere damage.

Finally, prior work indicates that self-perspective management is not reducible to generic executive demands but involves control processes that are specifically engaged when mental state representations conflict [97,98]. Similarly, working memory predicts ToM in different clinical populations but does not fully account for performance [86,99]. Together with evidence that ToM impairments after acquired brain injury are not fully explained by executive dysfunction [100–103] (for reviews see Aboulafia-Brakha et al. [104]; Wade et al. [105]), our findings support the interpretation that ToM-specific self-perspective conflict management is a distinct contributor to conversational success following RHD.

### Implications

Is ToM a required ability for successful conversation? One possibility is that much real-time interaction can be supported by relatively efficient forms of perspective tracking under low self-perspective conflict, with more flexible and cognitively demanding belief reasoning becoming important when the self and other perspectives diverge. This idea is compatible with dual-process accounts which distinguish between two belief reasoning systems: an efficient but limited one and a more flexible but resource-demanding one (see Westra & Nagel [29]; Butterfill & Apperly [34]). On this view, a structured collaborative task like Diapix may be one of the situations in which the more demanding form of ToM is particularly important because success requires explicit management of conflicting perspectives, a cognitively demanding task. An alternative view is that belief reasoning is engaged more pervasively during dialogue, including ordinary comprehension and production. Rubio-Fernandez and colleagues [78,79] propose that everyday conversation only works the way it does due to its reliance on belief reasoning ability (e.g. how do we foresee and prevent misunderstandings without reasoning with other’s beliefs?; cf. Heller & Brown-Schmidt [30,31]; Mazzarella et al. [106]). Our results do not fully adjudicate between these possibilities. What they do suggest is that accounts of cooperative dialogue that rely only on domain-general, non-mindreading processes are insufficient to explain this kind of interaction, because performance on the high self-perspective conflict condition explained unique variance in conversational success beyond broader cognitive measures (cf. Heyes [107]; Kissine [108]). Future work should test whether lower conflict forms of perspective tracking are sufficient for less structured conversational exchanges.

Overall, these findings reinforce the view that communication after RHD relies on social cognitive mechanisms, not just linguistic or general cognitive processes. Standard language and cognitive assessments early after stroke, which focus on motor-speech, language abilities, and visuospatial and executive skills, may miss the subtle conversation-relevant deficits stemming from impaired ToM. Incorporating socio-cognitive screening in clinical assessment early after RHD could help identify patients who need targeted support for communicative abilities [109]. Additionally, ToM-based interventions may represent one route to improve functional communication. Acknowledging differences in congenital versus acquired neurological conditions, evidence from autism interventions suggests that targeted ToM training can improve some communicative behaviors, underscoring the need for analogous work in stroke rehabilitation [110–112]. Initial rehabilitation studies targeting ToM in single cases of acquired brain injury (including RHD) suggest ToM abilities can improve [113,114], although effects on communication outcomes remain largely untested. Relatedly, emerging automated natural language processing approaches that quantify linguistic markers of ToM in spontaneous speech (e.g. mental state terms, referring expressions, pragmatic markers) could aid identification and monitoring of social-cognitive communication impairment in clinical settings [115].

### Limitations and Future Directions

This study advances what we know about RHD communication by isolating self ToM as a key contributor of conversational success while controlling for multiple demographic and cognitive factors, by using a structured yet natural conversation task that supports a quantitative measure of conversational success, by minimizing language confounds via a fully non-verbal ToM measure, and by testing individuals early post-stroke when neural reorganization and compensatory strategies are less likely to obscure disrupted mechanisms. At the same time, there were limitations. Although this is the only study to examine the relationship between conversation and ToM in early RHD, the sample size reflects the challenges of recruiting and testing medically vulnerable individuals in the acute phase of stroke (also see Kim et al. [116], n = 11) and the findings should be replicated in a larger sample. The conversation task emphasized explicit reasoning of the partner’s knowledge and structured turn-taking more than spontaneous everyday conversation. This structure may have increased the relevance of self-perspective management and underestimated ToM effects that emerge in less constrained interactions.

Finally, we observed an unexpected negative association between the self ToM control composite score and conversational success across several model specifications. When we decomposed this composite score into its constituent trial types (filler, memory-control, and true belief trials), the negative association attenuated and no single constituent measure was significantly associated with conversational success, while self ToM false belief reasoning remained a robust predictor. Because this control effect was unpredicted and could not be isolated to a specific task-processing component, it should be interpreted with caution. Further work with a larger sample will be needed to determine whether this association is reproducible and what processes might underlie it.

Future work can extend these findings in several directions. Fine-grained analyses like exchange structure analysis [117] (also see, Kilov et al. [118]) could identify where conversation breakdowns occur and which communication strategies (e.g. requests for clarification, information-seeking questions, repairs) covary with ToM. For example, based on our interpretation that belief reasoning is involved in coordinating shared understanding during collaborative interaction, one possibility is that individuals with poorer ToM will request less information from a conversation partner and ask fewer clarifications when the partner’s utterances are ambiguous. Testing whether these behaviors mediate the relationship between ToM and conversational success would help specify the mechanism through which ToM supports communication. Longitudinal designs will also be important for determining whether changes in ToM predict changes in conversational success over recovery. If the relationship between ToM and conversation is stable over time, that would provide stronger evidence for a mechanistic link between disrupted ToM and functional communication outcomes.

## Conclusions

In adults tested early after RHD, conversational success was uniquely linked to the ability to resist interference from one’s own privileged knowledge to explicitly reason about a partner’s perspective when the two conflict. This relationship remained after accounting for demographics and broader cognitive abilities, whereas inferring what another person knows or believes under lower perspective conflict (other ToM) showed no significant association. These findings suggest that social-cognitive mechanisms, specifically self-perspective conflict management during belief reasoning, are a distinct contributor to collaborative communication after RHD. They motivate future work to test when different components of ToM matter most for dialogue and to evaluate whether socio-cognitive assessment and intervention can improve real-world communication after stroke.

## Data and code availability statement

The data supporting the findings of this study are subject to HIPAA regulations. Analytical code and non-identifying data are available at Zenodo repository: https://doi.org/10.5281/zenodo.23568794

## Acknowledgements

Thanks to our participants, their care partners and the stroke units at the Texas Medical Center; Danielle Brown (DB), Emilia Cichoki (EC), Kennedy Guess (KG) and Somya Mittal (SM) for data collection; Gerri Esquivel (GE) and Junha Lee (JL) for transcription assistance; Kesha Pugalenthi and the Texas Advanced Computing Center at UT Austin for access to the lonestar6 system for Whisper-large-V3 model for automated transcription; statistical consultant John Magnotti; and lesion tracing consultant Chris Hamilton (CH).

## Author contributions

Andrea Suazo (AS): Investigation, Formal Analysis, Visualization, Writing – Original Draft. Margaret Blake: Conceptualization, Funding acquisition. Tatiana T. Schnur: Conceptualization, Funding Acquisition, Methodology, Project administration, Supervision, Writing – Original Draft.

## Funding

This project was supported by the NIH-NIDCD award #R01DC019828 to UTHealth Houston.

## Additional Information

The authors report no competing interest.

## Notes

### Competing Interest Statement

The authors have declared no competing interest.

### Summary of Updates

We addressed reviewer concerns regarding the anatomical heterogeneity introduced by two participants with predominantly right cerebellar strokes. These participants met the original study eligibility criteria and were therefore included in the original analyses. However, given the reviewer concerns and the predominantly right cerebral hemisphere composition of the sample, we revised the primary analytic cohort to exclude these two cases. We updated the participant characteristics, lesion distribution figure, figures, tables, statistical analyses, and Supplementary Information accordingly. Importantly, this change did not alter the central conclusion: self ToM false belief performance remained significantly associated with conversational success in the revised primary model (std. beta = .42, p = .023) and across the sensitivity analyses (std. betas = .44 to .64, ps = .004 to .017). We also addressed a question regarding overall lesion burden by conducting an additional targeted sensitivity analysis incorporating lesion volume. In the subset with available lesion volume data (N = 24), self-ToM was a significant predictor of conversational success after adjustment for lesion volume (std. beta = .48, p = .033), whereas lesion volume itself was not significantly associated with conversational success (std. beta = -.28, p = .223). This analysis is reported in the Supplementary Information.

