## Supplementary Material for "Theory of mind predicts conversational success in early right hemisphere stroke recovery"

### Supplementary Methods

#### Image Acquisition and Analysis

To determine the distribution of acute brain damage in the patient cohort, we acquired neuroimaging scans as a part of standard clinical care. MRI was contraindicated for 6 participants. For the remaining 25 participants, we acquired diffusion weighted, apparent diffusion coefficient, and high-resolution structural scans (T1 and T2 FLAIR) along the axial direction with 4–5 mm slice thickness. The voxel sizes of diffusion-weighted and structural images were approximately 1×1×4.5 mm, and 0.5 × 0.5 × 4.5 mm, respectively.

To demarcate lesions, we first registered the diffusion weighted images with the high-resolution structural images (T1 or T2) using AFNI (https://afni.nimh.nih.gov/). Lesions were demarcated directly on the diffusion weighted images (by CH), using ITK-snap (http://www.itksnap.org/pmwiki/pmwiki.php) with reference to apparent diffusion coefficient and T2 FLAIR images. Next, we normalized the individual structural images to the Colin-27 template in Montreal Neurological Institute (MNI) space using ANTS registration (http://stnava.github.io/ANTs/) [1]. We used the corresponding affine parameter and diffeomorphic maps to warp individual masks to the MNI space [2,3].

We created a lesion overlap map and determined regions of highest overlap according to the AAL atlas in MRIcro. Lesion volume was calculated using the fslstats function from the fslr package in R [4].

#### Transcription and Interrater Reliability

We transcribed Diapix conversation using an artificial intelligence model for automatic speech transcription (whisper large-v3 model) [5]. A research team member (AS, GE, JL) manually checked transcripts for word identification and utterance boundaries, adjusting when necessary. Utterance boundaries were identified using grammatical clauses outlined by Salt Software [6]. We established interrater reliability on 10 randomly selected RHD participant conversations using point-to-point agreement based on total number of words, word identification and utterance boundaries. Research team members reached an average of 96.5% agreement on word identification (range 93.8% –98.8%), 97.9% agreement on total number of words (range 92.4% –99.9%) and 95.8% agreement on segmentation of utterances (range 90.5% –100%).

**Supplementary Figures**

**Supplementary Figure S1. Study procedure flowchart.** Flowchart illustrating the testing battery and protocol sequence. Theory of Mind (ToM) assessments were administered in a fixed sequence (Other ToM followed by Self ToM); however, the administration order of remaining baseline cognitive and language assessments varied based on patient constraints.

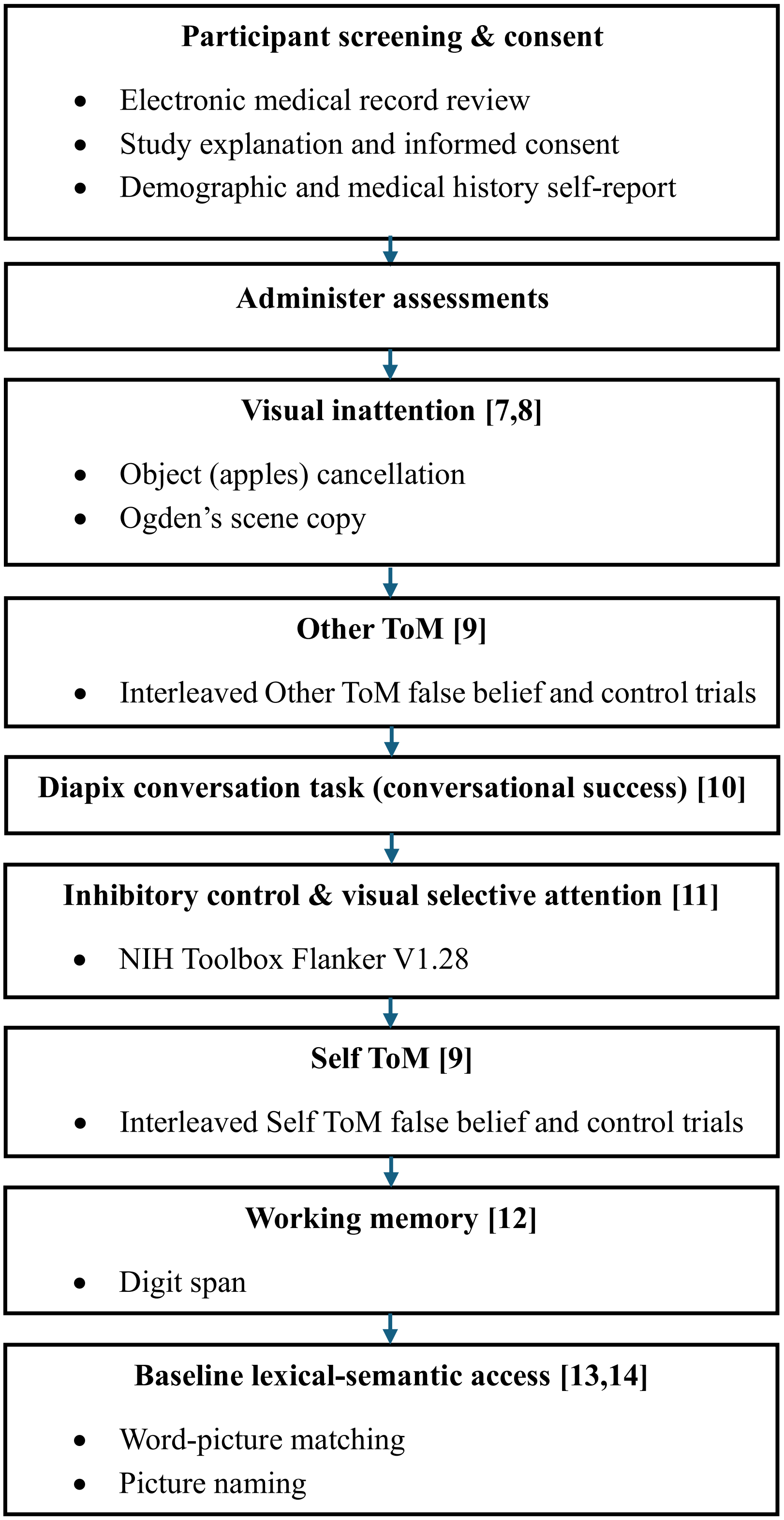

**Supplementary Figure S2. Baseline cognitive performance in early post-stroke right-hemisphere damage (RHD).** Violin plots illustrate performance distributions, where circles represent individual participant *z*-scores and inset boxplots denote the median and interquartile range. The dashed line marks $z$ = 0 (reference mean). Visual inattention *z*-scores are standardized to the RHD sample (n = 31), inhibitory control/ attention to the demographically matched NIH Toolbox normative sample (n = 1,038) [11], and working memory to the control sample (n = 16).

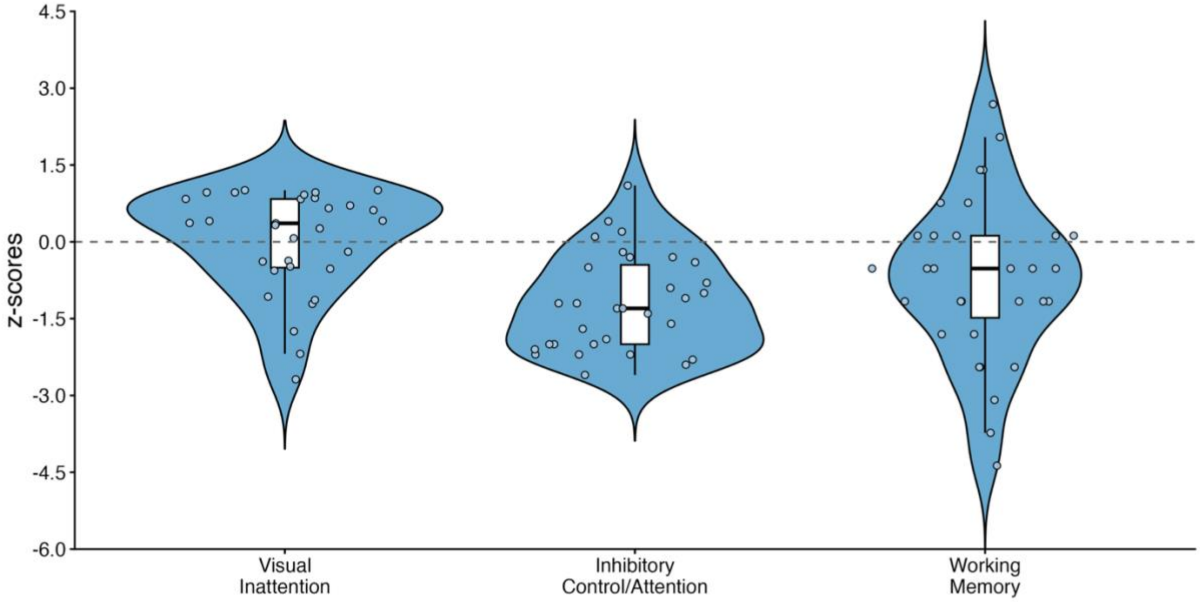

**Supplementary Figure S3. Baseline expressive and receptive lexical-semantic access in early post-stroke right-hemisphere damage (RHD).** Violin plots illustrate performance distributions across RHD participants (n=29), where circles represent individual participant *z*-scores and inset boxplots denote the median and interquartile range. The dashed line marks $z = 0$ (reference mean). Word-picture matching *z*-scores are referenced to the full control sample (n = 16), and picture naming to a reduced control sample (n = 15; one control participant did not complete the task).

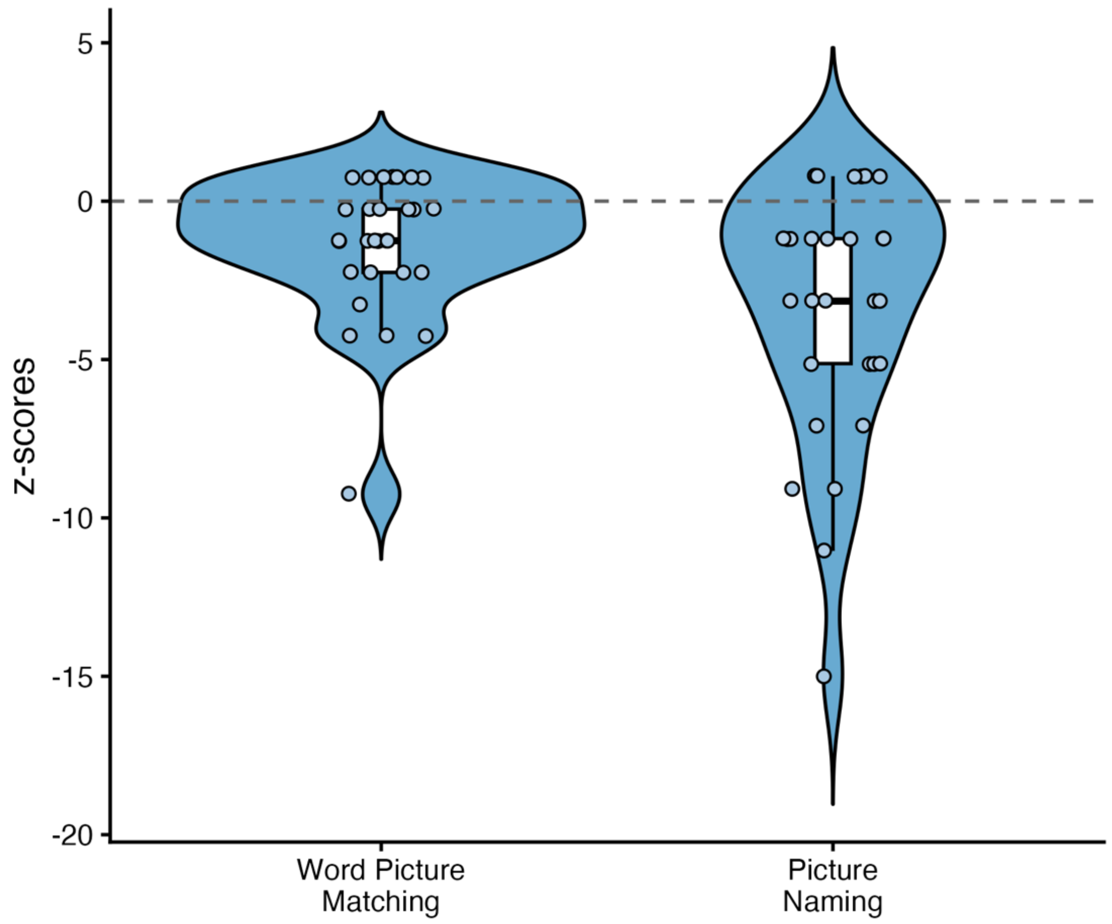

**Supplementary Figure S4. Extended correlation matrix across demographics, lesion size, baseline cognition, language, Theory of Mind (ToM), and conversational success in early post-stroke right-hemisphere damage (RHD).** Matrix cells report pairwise Pearson's r correlation values across the RHD group (n = 30; n = 28 for lexical-semantic language tasks; n = 24 for lesion size [$cm^3$]). Colored cells indicate significant correlations ($p$ < .05). Circle size and color intensity indicate the magnitude and direction of the association (see color bar).

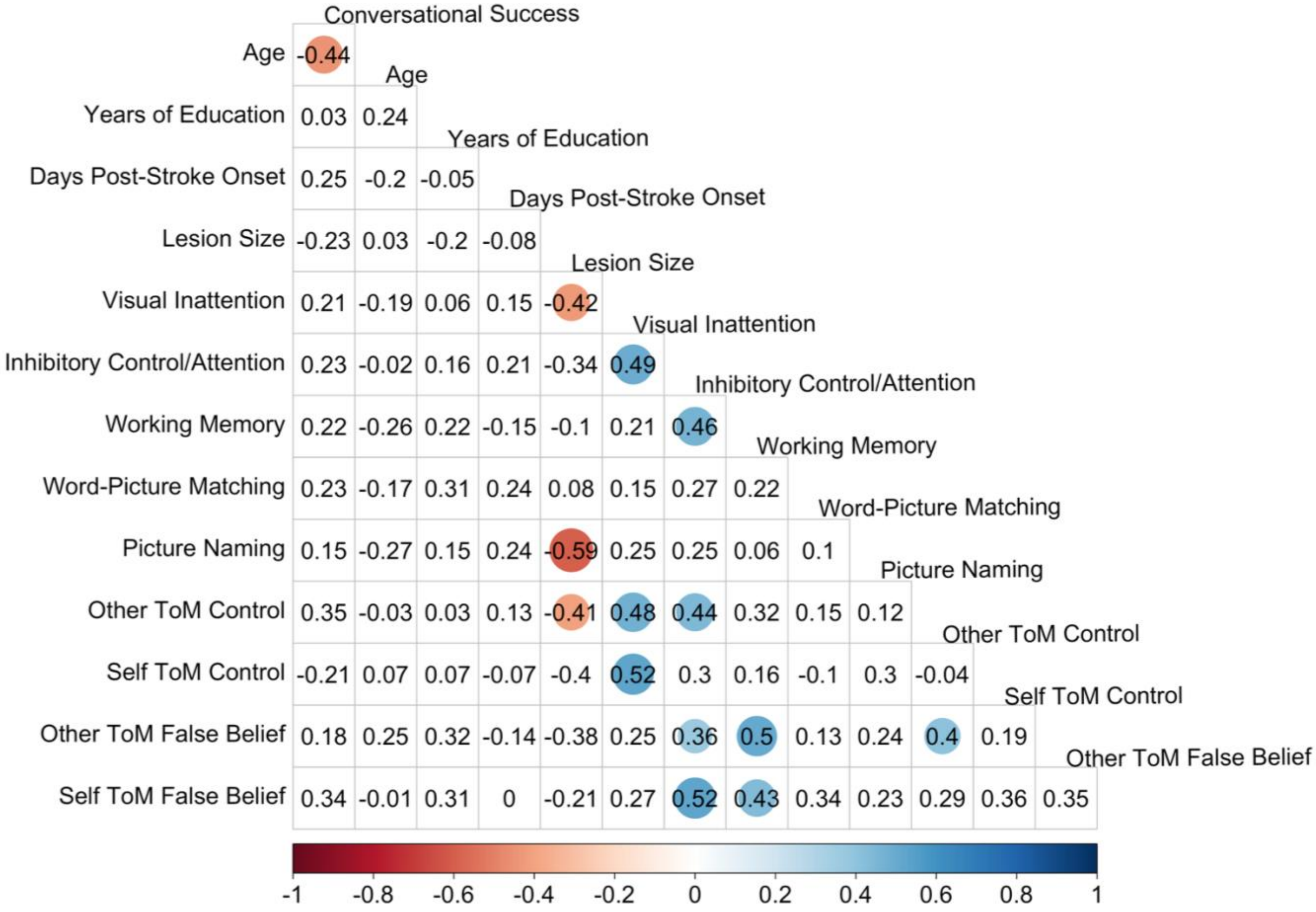

**Supplementary Figure S5. Task accuracy across Theory of Mind (ToM) false belief and control conditions in early post-stroke right hemisphere damage (RHD) and healthy controls.** Grouped boxplots display percentage accuracy across the four ToM conditions for the RHD (n = 31) and control (n = 16) groups. Center lines indicate the median, boxes represent the interquartile range, and whiskers indicate the overall range. Overlaid symbols represent individual participants (RHD, blue circles; controls, red triangles).

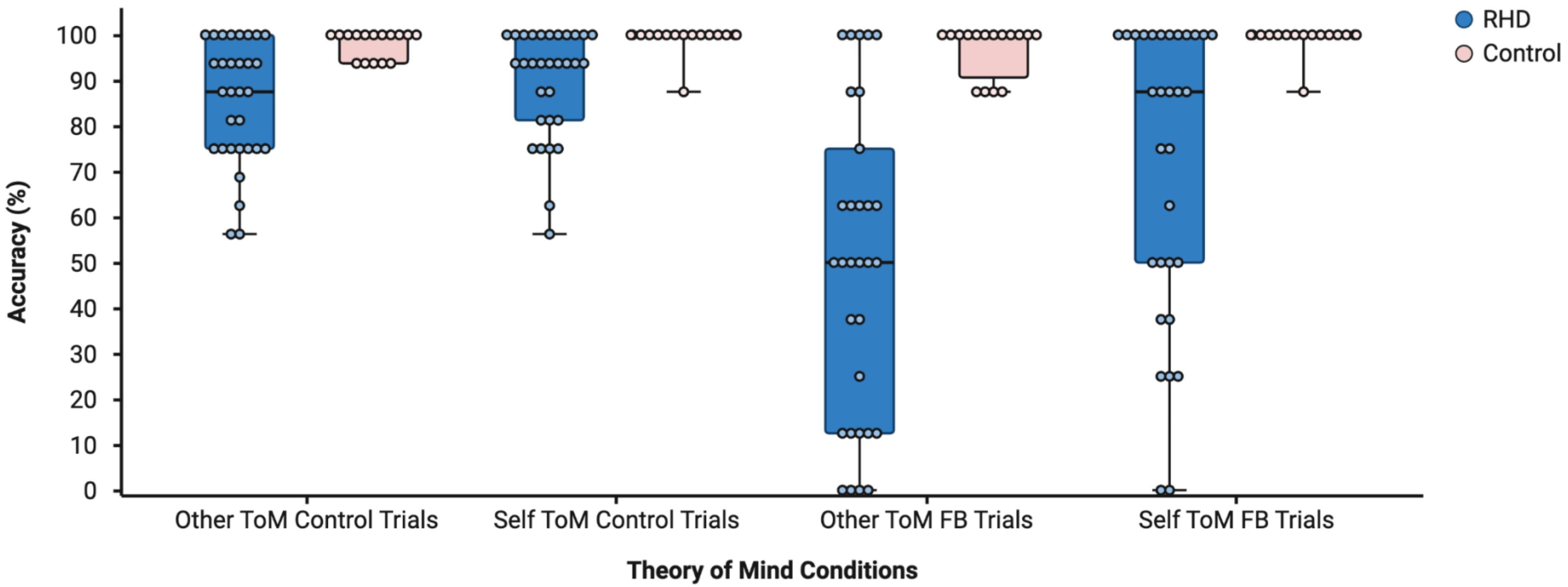

**Supplementary Figure S6. Conversational success in early post-stroke right-hemisphere damage (RHD) and healthy controls.** Boxplots show conversational success (percentage of total differences identified; 12 possible) for the RHD (n = 31) and control (n = 16) groups. Center lines indicate the median, boxes represent the interquartile range, and whiskers indicate the overall range. Overlaid symbols represent individual participants (RHD, blue circles; controls, red triangles).

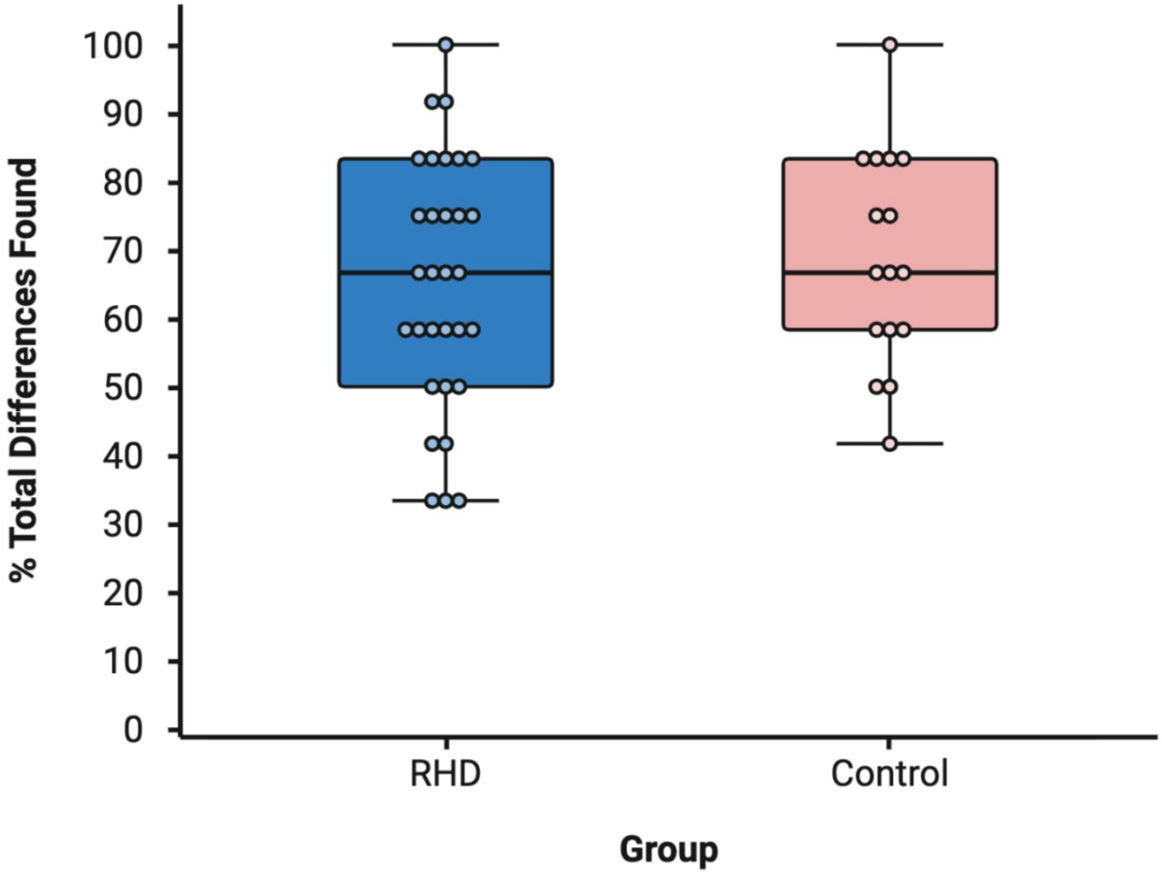

**Supplementary Figure S7. Sensitivity regression models 1 and 2 predicting conversational success in early post-stroke right-hemisphere damage (RHD; n = 30).** Standardized fixed-effect estimates (β) with 95% confidence intervals from regression models predicting conversational success. (A) Sensitivity model 1 (cognition-adjusted) incorporating Theory of Mind (ToM) predictor composites with baseline cognitive covariates (visual inattention, inhibitory control/attention, working memory). (B) Sensitivity model 2 (demographics-adjusted) incorporating ToM predictors with demographic and clinical covariates (age, years of education, days post-stroke onset). Dark blue bars denote significant predictions ($p < .05$); light blue bars denote non-significant predictors ($p > .05$). Predictors ordered by standardized beta value.

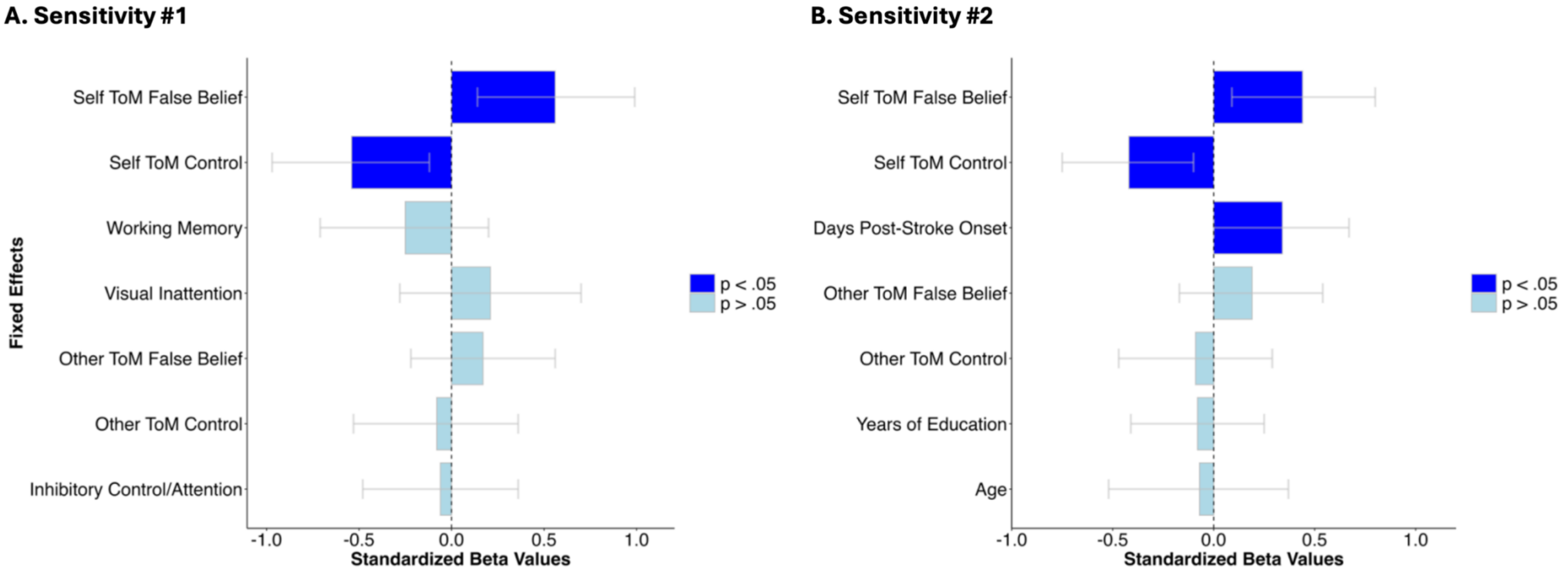

**Supplementary Figure S8. Adjusted partial regression plots relating conversational success and Theory of Mind (ToM) performance in early post-stroke right-hemisphere damage (RHD; n = 30).** Partial regression plots derived from the fully adjusted (Sensitivity 3) model. Each panel displays the partial association between *z*-scored conversational success and *z*-scored ToM performance after adjusting for demographic and baseline cognitive covariates. Blue circles represent individual participants; solid lines show model fits; shaded bands indicate 95% confidence intervals.

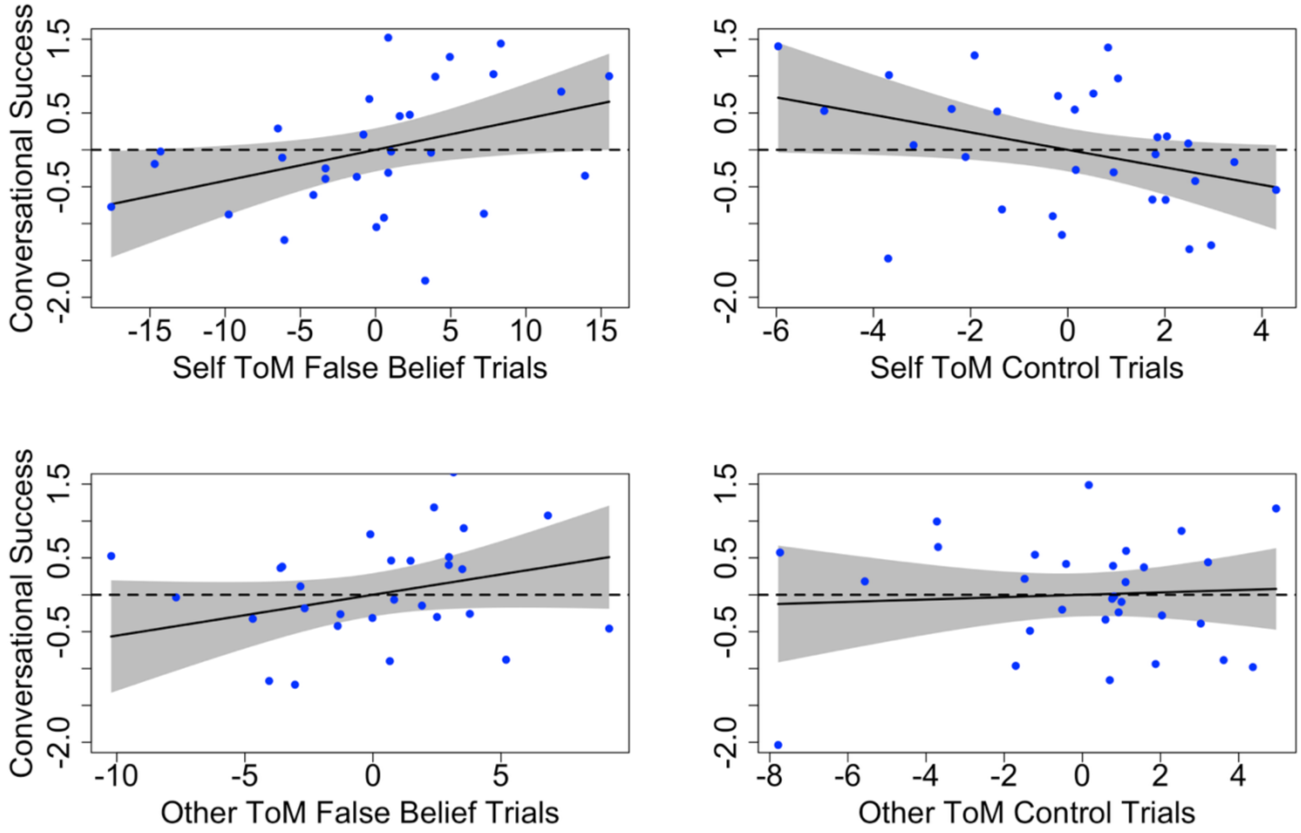

**Supplementary Figure S9. Primary model predicting conversational success with decomposed Theory of Mind (ToM) control trials (n = 30).** Standardized fixed-effect estimates (β) with 95% confidence intervals from the exploratory trial-type decomposition model. Dark blue bars indicate significant predictions (*p* < .05). Light blue bars indicate non-significant predictors (p >= .05). Predictors ordered by standardized beta value.

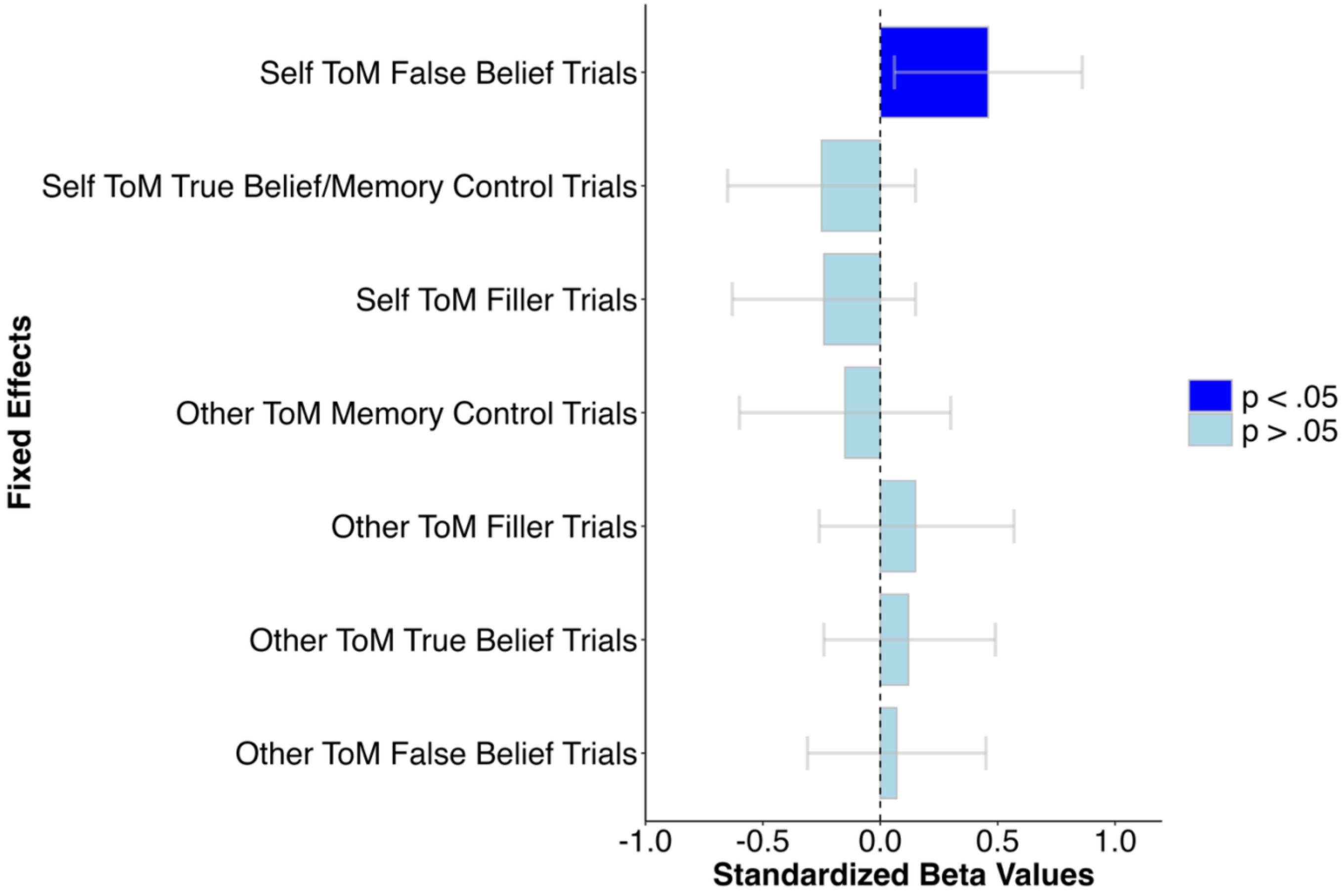

**Supplementary Tables**

**Supplementary Table S1. Baseline cognitive and lexical-semantic access task performance in early post-stroke right-hemisphere (RHD) and healthy controls.**

| Task | RHD (n = 31) | | Control (n = 16) | |
|---|---|---|---|---|
| | Mean (SD) | Range | Mean (SD) | Range |
| Visual inattention (accuracy) | 92% (8%) | 71% - 100% | N/A | N/A |
| Inhibitory control & visual selective attention (NIH Toolbox flanker t-score) | 38 (9) | 24 - 61 | 54 (7) | 43 - 66 |
| Working memory (digit span raw score) | 11 (2) | 5 – 16 | 12 (2) | 8 – 14 |
| Word-picture matching accuracy | 96% (4%) | 80% - 100% | 99% (2%) | 94% - 100% |
| Picture naming accuracy | 93% (7%) | 73% - 100% | 99% (2%) | 96% - 100% |

Note: Values represent group mean (SD) and overall range. Visual inattention reflects composite accuracy from the Apples Cancellation [7] and scene copy tasks [8]. Inhibitory control/visual selective attention reflects the NIH Toolbox Flanker Inhibitory Control and Attention Test T-score (V1.28) [11]. Working memory reflects the Repeatable Battery for the Assessment of Neuropsychological Status (RBANS) digit span (raw score) [12]. A subset of 29 RHD participants completed the word-picture matching [13] and picture naming tasks [14]. Visual inattention was not administered to control participants due to ceiling performance. Ten controls completed the Flanker task (administered in person only); 15 controls completed the picture naming task.

**Supplementary Table S2. Primary regression model predicting conversational success with decomposed Theory of Mind (ToM) control trials in early post-stroke right-hemisphere damage (RHD; n = 30).**

| | **Decomposed Primary Model** | | |
|---|---|---|---|
| **Fixed Effects** | **std. β** | **std. 95% CI** | ***p*** |
| Other ToM Filler Trials | 0.15 | -0.26, 0.57 | .450 |
| Other ToM Memory Control Trials | -0.15 | -0.60, 0.30 | .505 |
| Other ToM True Belief Trials | 0.12 | -0.24, 0.49 | .484 |
| Self ToM Filler Trials | -0.24 | -0.63, 0.15 | .211 |
| Self ToM True Belief/Memory Control Trials | -0.25 | -0.65, 0.15 | .203 |
| Other ToM False Belief Trials | 0.07 | -0.31, 0.45 | .689 |
| Self ToM False Belief Trials | 0.46 | 0.06, 0.86 | .027* |
| **Random Effects** | **RV (σ2)** | **RIV (τ00)** | **ICC** |
| Conversation Partner (n=3) | 0.87 | 0.24 | 0.22 |

Note: Values represent standardized regression estimates (β), 95% confidence intervals (CI), and associated *p*-values. Statistically significant effects ($p < .05$) are marked with an asterisk (*). std. = standardized; std. = standardized, CI = confidence interval; ICC = intraclass correlation coefficient; RV = residual variance ($\sigma^2$); RIV = random intercept variance ($\tau_{00}$); ToM = Theory of Mind.

**Supplementary Table S3. Targeted lesion-volume sensitivity analysis predicting conversational success in early post-stroke right hemisphere damage**

| | Primary Model (N=24) | | | Primary Model + Lesion Volume (N=24) | | |
|---|---|---|---|---|---|---|
| **Fixed Effects** | **std. β** | **std. 95% CI** | ***p*** | **std. β** | **std. 95% CI** | ***p*** |
| Other ToM Control Trials | 0.21 | -0.26, 0.68 | .366 | 0.11 | -0.39, 0.61 | .648 |
| Self ToM Control Trials | -0.42 | -0.85, 0.01 | .057 | -0.53 | -1.00, -0.06 | .029* |
| Other ToM False Belief Trials | -0.01 | -0.45, 0.42 | .952 | -0.05 | -0.49, 0.38 | .801 |
| Self ToM False Belief Trials | 0.44 | 0.00, 0.88 | .050* | 0.48 | 0.04, 0.92 | .033* |
| Lesion Volume | | | | -0.28 | -0.74, 0.18 | .223 |
| **Random Effects** | **RV (σ2)** | **RIV (τ00)** | **ICC** | **RV (σ2)** | **RIV (τ00)** | **ICC** |
| Conversation Partner (n=3) | 0.86 | 0.11 | 0.11 | 0.84 | 0.09 | 0.10 |

Note: Values represent standardized regression estimates (β), 95% confidence intervals (CI), and associated *p*-values. Statistically significant effects ($p < .05$) are marked with an asterisk (*). std. = standardized; std. = standardized, CI = confidence interval; ICC = intraclass correlation coefficient; RV = residual variance ($\sigma^2$); RIV = random intercept variance ($\tau_{00}$); ToM = Theory of Mind.
